# Mitochondrial Phenotypes Distinguish Pathogenic MFN2 Mutations by Pooled Functional Genomics Screen

**DOI:** 10.1101/2021.03.12.434746

**Authors:** Alex L. Yenkin, John C. Bramley, Colin L. Kremitzki, Jason E. Waligorski, Mariel J. Liebeskind, Xinyuan E. Xu, Maria A. Vakaki, Vinay D. Chandrasekaran, Robi D. Mitra, Jeffrey D. Milbrandt, William J. Buchser

## Abstract

Most human genetic variation is classified as VUS - variants of uncertain significance. While advances in genome editing have allowed innovation in pooled screening platforms, many screens deal with relatively simple readouts (viability, fluorescence) and cannot identify the complex cellular phenotypes that underlie most human diseases. In this paper, we present a generalizable *functional genomics* platform that combines high-content imaging, machine learning, and microraft isolation in a new method termed “Raft-Seq”. We highlight the efficacy of our platform by showing its ability to distinguish pathogenic point mutations of the mitochondrial regulator *MFN2*, even when the cellular phenotype is subtle. We also show that our platform achieves its efficacy using multiple cellular features, which can be configured on-the-fly. Raft-Seq enables a new way to perform pooled screening on sets of mutations in biologically relevant cells, with the ability to physically capture any cell with a perturbed phenotype and expand it clonally, directly from the primary screen.

**Graphical Abstract:** 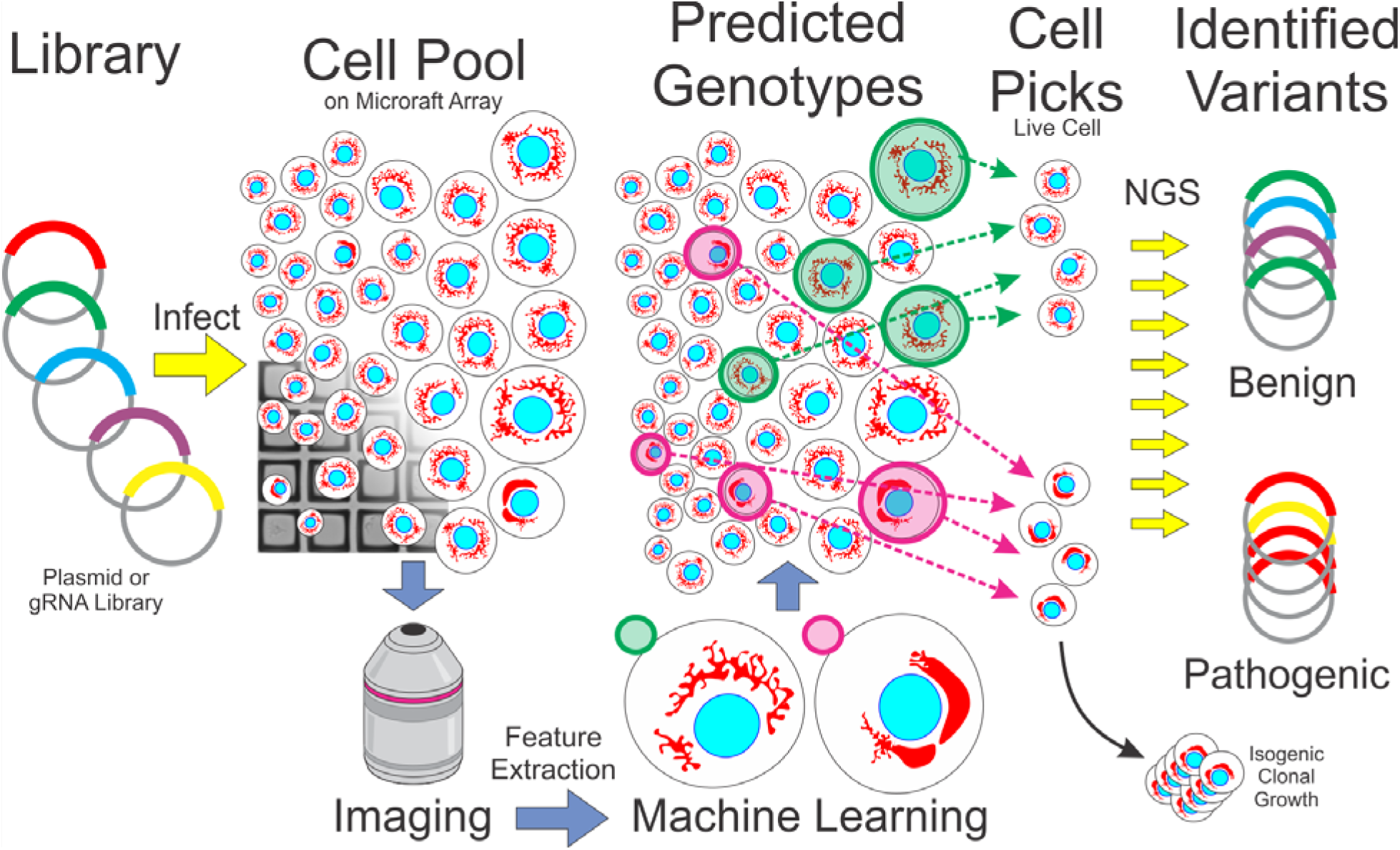

Here, we address the need to evaluate the impact of numerous genetic variants. This manuscript depicts the methods of using machine learning on a biologically relevant phenotype to predict specific point mutations, followed by physically capturing those mutated cells.

## Background

The explosion of functional genomics in the past decade ^1^ has enabled a massive shift in the study of the underlying genetics of human pathology. Even so, it is difficult to connect specific genetic mutations to disrupted cellular phenotypes, necessitating a detailed phenotyping-based *functional genomics* platform that can both screen large numbers of genetic perturbations/variants—so called multiplexed assays of variant effects (MAVE) ^2^—and work alongside new methods of generating variant libraries, such as deep mutational scanning (DMS) ^3^.

Genetic perturbation screens have generally relied on simple phenotypes conducive to enrichment analysis ^4^, Fluorescence-Assisted Cell Sorting ^5–7^, or phenotypes measurable by sequencing ^8–10^. For more complex phenotyping, high-content imaging/screening (HCI/HCS) ^11–15^ is performed in an arrayed format for genetic perturbation screens ^16–21^, but these are difficult to scale and infeasible for combinatorial screens. There are platforms that use an imaging-based approach in pooled genetic perturbation screens, and have been demonstrated on relatively simple phenotypes ^22–, 24^ or on precise phenotypes that were known in advance ^17, 25^.

Although some platforms do isolate individual cells ^26^, most genetic perturbation screens use a population-level measurement as their endpoint. The main challenge for such screens that operate on the single-cell level is mapping the perturbation back to each cell post sequencing. For screens where the phenotype is measurable through sequencing, the perturbagen can be found simultaneously with the phenotype with single-cell resolution^8, 10^, and newer methods have incorporated more advanced sequencing techniques to find additional data, such as surface protein presence^10^. In imaging, a cell’s position within a microscopic field of view is the unique piece of identifying information used to map phenotype to perturbation. Some platforms use *in situ* sequencing ^27, 28^ to generate sequencing results that contain positional data to a specific cell ^17, 29^. Other platforms use a digital micromirror device to photoactivate endogenous fluorophores in specific cells which are then put through FACS-seq ^23, 25, 30^. However, a platform that can be integrated into existing next-generation sequencing (NGS) pipelines would be more accessible and robust.

In this paper, we present Raft-Seq, a pooled screening platform that predicts individual cell perturbations from high-content imaging and machine learning. Raft-Seq improves on other platforms in several important ways: 1, it uses a microraft plate ^31–33^ for context-aware isolation of identified cells; 2, it can use vital dyes or stains, so no genetic modification of the cell is required other than the perturbation itself; 3, it is largely phenotype-agnostic, needing only knowledge of the broad physiology beforehand for stain selection and initial feature filtering; 4, it uses machine learning to identify perturbed cells, allowing the identification of complex cell-autonomous phenotypes; 5, it selects cells with high viability and clonability, directly from the primary screen.

Here, we use the Raft-Seq platform to examine the neurologically relevant mutations in the *MFN2* gene, which protects against cellular stress from damaged mitochondria by regulating mitochondrial fission and fusion ^34^. Clinically relevant *MFN2* mutations (pathogenic variants) primarily result in Charcot-Marie-Tooth Disease (CMT), the most common inherited neuromuscular disorder characterized by peripheral neuropathy with impairment of the central nervous system ^35–38^. We find that the phenotype caused by pathogenic *MFN2* variants is distinct, but the difference is not adequately described by a single measurement/feature, necessitating a more complex feature analysis. Following the findings from the single perturbation experiments, we targeted a gRNA library across the *MFN2* coding region to identify anomalous phenotypes caused by these mutations.

## Results

### MFN2 Mutants

We generated cell lines that each contain *MFN2* cDNA with single point mutations through lentiviral infection. The mutants studied consisted of six clinically relevant pathogenic *MFN2* variants ^39^ and five control variants annotated as “benign” from ClinVar ^40^ (**Figure 1a**). Twelve lentiviral transfer plasmids—one plasmid for each mutated cDNA and one for wild type cDNA—were used to package lentivirus that was infected into U2OS to create twelve cell lines.

**Figure 1.**
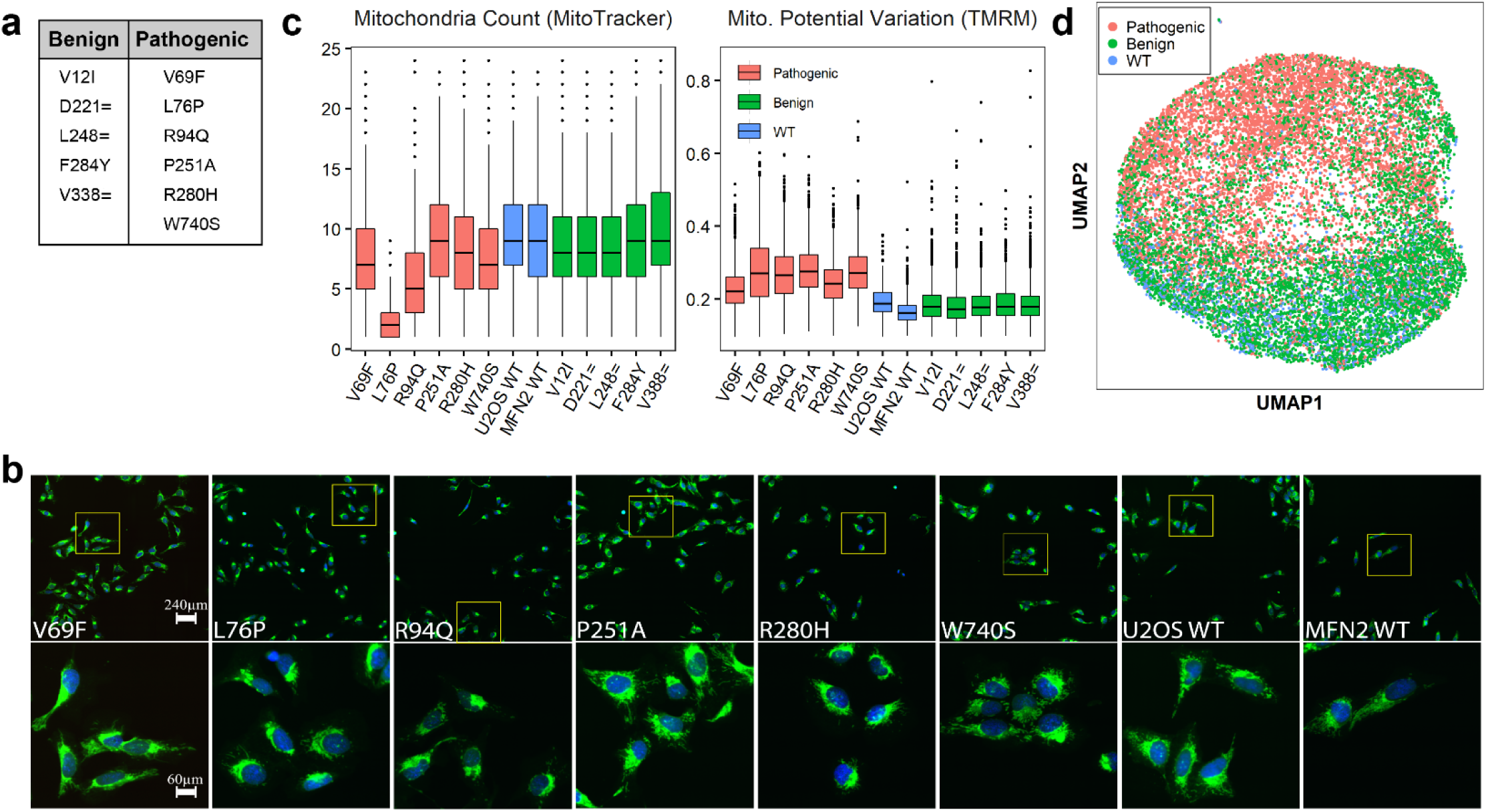
Isogenic *MFN2* Mutant Cell Lines Characterization: **a.** Table of the mutants used. **b.** Confocal images of mutant cells at two magnifications (blue = nuclear staining by Hoechst, green = mitochondria staining by MitoTracker). Subtle mitochondrial differences can be observed amongst the mutant populations. Known pathogenic mutants display peri-nuclear aggregation of mitochondria and a lack of mitochondrial spreading. In comparison, wild-type cells show ample mitochondrial spreading. **c.** Box plots of two features: Mitochondrial count (based on MitoTracker staining) and coefficient of variation in mitochondrial potential (TMRM intensity). Histograms are colored by genotype class (Benign/Pathogenic/WT). **d.** Scatter plot of UMAP reduction using a set of 26 features measured with TMRM or MitoTracker staining. Each dot represents a single cell, colored by its genotype class (WT/benign/pathogenic).

To investigate the phenotypes caused by the *MFN2* variants, we assessed mitochondria and mitochondrial membrane potential using MitoTracker and TMRM, respectively ^41^. The cells were imaged using a high-throughput fluorescent confocal microscope and intracellular features were measured using Cytiva’s IN Carta analysis software. A visual comparison of the *MFN2* cell lines demonstrates subtle phenotypic differences among them (**Figure 1b**), specifically, perinuclear aggregation of mitochondria in the cells containing the pathogenic mutations. Box plots for a subset of the features are shown in **Figure 1c.** The mean mitochondrial puncta count in the wild-type and benign cell lines is consistent, while the values for the pathogenic cell lines were lower and more variable. A higher average variation in TMRM intensity within a single cell was also observed in the pathogenic mutants, all suggesting mitochondrial aggregation and disrupted mitochondrial membrane potential. Additional feature histograms are shown in **Figure S1**. Although there are significant differences in the population of mutants as a whole, there is no single feature that can separate more than a small fraction of the cells.

We next tested whether multiple features could distinguish the pathogenic-variant cells. UMAP dimensionality reduction was performed on the full set of cellular image-based features (**Figure 1d**). Most pathogenic mutant cells are separable from wild type/benign cells in the dimensionality reduction, though some mutants (L76P and R94Q) are more distinct (**Figure S2**), which is consistent with the individual feature data. The UMAP reduction shows that using data from multiple features may more reliably distinguish cells with pathogenic mutants. Thus, to screen individual cells based on subtle phenotypes caused by these pathogenic mutations, we designed an HCS platform with machine learning capabilities that can use information from multiple image features.

### Experimental Workflow

The overall workflow of Raft-Seq can be summarized in five steps: perturbation, imaging, model building, isolation, and sequencing (**Figure 2a**). In the first step, we introduce a genetic perturbation, either endogenous (e.g., CRISPR gRNA library) or exogenous (e.g., plasmid overexpression library with mutagenesis). The cells are then seeded onto a microraft plate, stained, and imaged on a high-throughput confocal microscope. From the cell-feature matrix obtained from feature extraction, we generate many supervised learning models, used to predict whether there is a discernible perturbation in each cell. Several different model types are employed (**Methods: Machine Learning and Model Generation**), which are then evaluated and deployed to determine cells to be isolated. Isolation is performed using the Cell MicroSystems Air Instrument, where rafts containing cells of interest are automatically transferred with a magnet to a well of a 96-well plate. The cells on the isolated rafts are then genotyped to determine the perturbation present. The genotyping pipeline is performed on single cells, not pooled genetic material.

**Figure 2.**
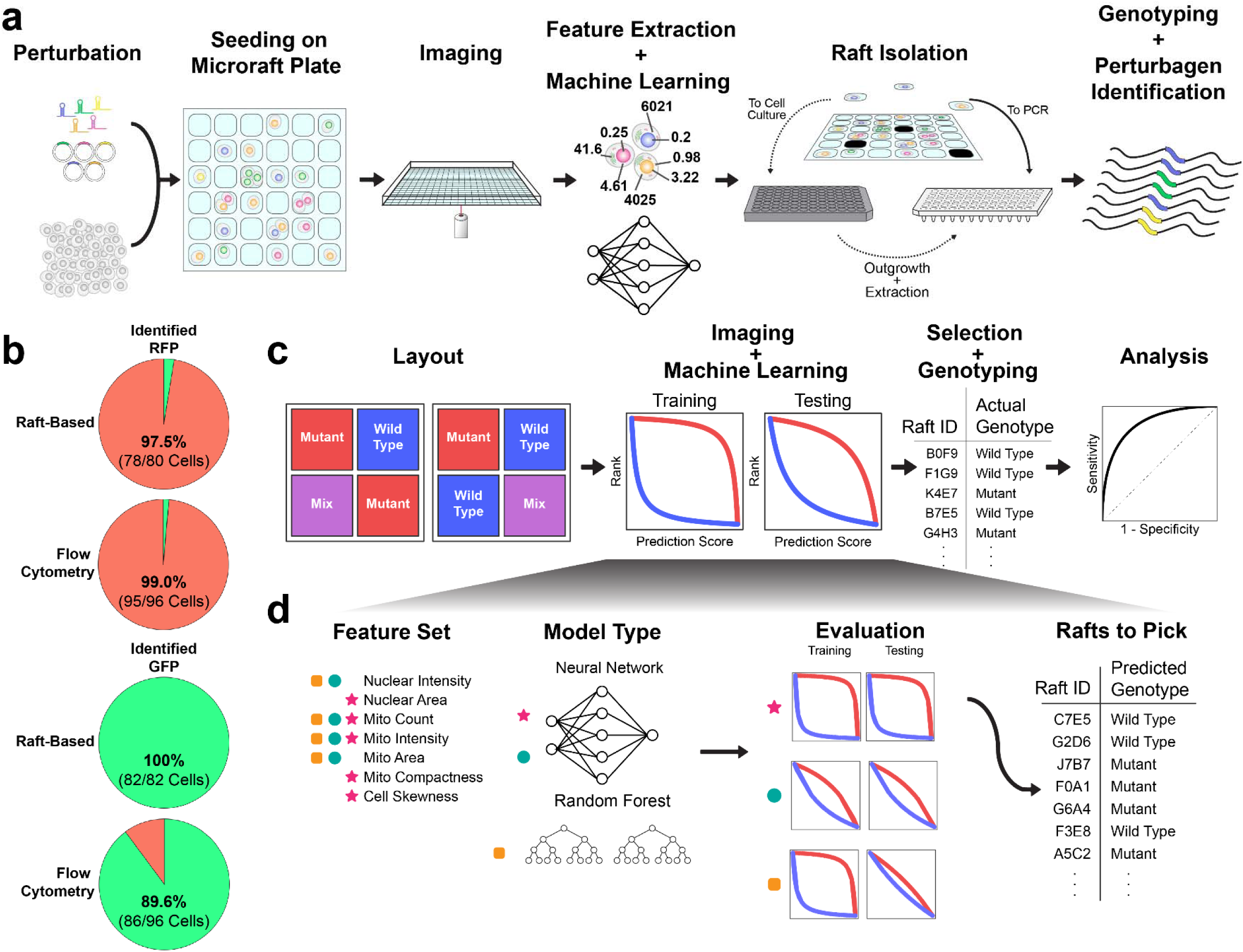
Raft-Seq Experimental Workflow. **a.** A perturbagen library, primarily a lentiviral gRNA or plasmid overexpression library, is introduced to cells, which are then seeded onto a microraft plate. The plate is imaged, and cell feature data are then extracted from the resulting images and used to build machine learning models. The trained model then selects rafts to either be isolated into a PCR plate for immediate analysis or a tissue culture plate for clonal expansion. Following isolation, the cells are genotyped. **b.** Pie charts showing the composition of cells identified as either expressing GFP or RFP, after being separated by either the raft-based approach or flow cytometry. The color represents the expected appearance of the cells based on genotype (green = contains RFP gRNA, red = contains GFP gRNA), and the fractions/percentages are the amount in each group that is correctly identified. **c.** Performance evaluation workflow. The blue and red wells represent the labeled cell populations and the purple well represents the unlabeled population that is a mixture between the two. Following imaging, models are trained and tested on the labeled wells, and the one with the clearest delineation between the two classes is selected. That model is used to select rafts that are then isolated and single cell genotyped. The comparison between the prediction scores and the true genotype class generates a receiver operator characteristic curve to evaluate model efficacy. **d.** Modeling workflow. Each model comprises a feature set and machine learning algorithm, combinations of which are represented here with different symbols. Each model is evaluated with training and testing cell data, and the model that shows the largest separation between classes and is robust on the testing data is used to generate a list of cells (rafts) to pick, along with their predicted genotype.

As a preliminary test of raft-based imaging and isolation, we performed Raft-Seq on cells with dual genome-encoded RFP and GFP that had been given a gRNA to knock-out either RFP or GFP. These cells were plated on a microraft plate as a mixture and were imaged. Cells that were given the guide targeting RFP would be expected to appear green, since GFP would continue to be expressed and vice-versa. Using fluorescent marker intensity, we predicted the guide presence, isolated those cells, performed single-cell DNA sequencing to look for the edited genomic targets, and determined the true genotype. The correspondence between the predicted guide and the true guide was near perfect (Accuracy = 98.8%, n = 162) and comparable to a flow-based single-cell sequencing method run in parallel (**Figure 1b** and **Supplemental Methods**).

### Identifying Subtle Mitochondrial Phenotypes in a Mixed-Variant Pool

To validate the efficacy of Raft-Seq in a more complex screen, we attempted to separate a mixture of wild-type and mutant cells by reproducibly predicting a given cell’s *genotype* based on its *phenotype*, as determined by a set of features extracted from imaging data (with no fluorescent reporters). For this experiment, we separated cells containing pathogenic *MFN2* mutations from those containing the wild-type *MFN2* cDNA.

The workflow of this validation experiment is shown in **Figure 2c**. The goal was to see if we could identify any of four pathogenic mutants (L76P, R94Q, P251A, R280H) from WT in an admixture of those cells. In the different wells of the microraft plate, we plated either 1) *MFN2* WT cells, 2) a mixture of the 4 *MFN2* mutants (1:1:1:1), 3) a mixture of the wild-type and pathogenic cells at a wild-type:pathogenic ratio of 90:10, and 4) a similar mixture at a ratio of 50:50. We distinguished between the first two groups and the last two by calling the former (1,2) “labeled” and the latter (3,4) “unlabeled” to clarify for which cell populations the genotype class was known prior to imaging and analysis. Following application of a nuclear stain, a plasma membrane stain, and a mitochondrial stain (**Methods: Staining**), the cells were imaged and their features computed. Next, we designed classification models that best distinguished the labeled populations (*MFN2* WT vs. *MFN2* mutants). General modeling workflow is shown in **Figure 2d**. We used a variety of model algorithms (types) and cell features sets, preferentially choosing features that correlated with genotype class. For model selection, we trained each model on-the-fly using plate-spanning sets of feature data. For each set of predictions for each model, we graphed the ranked prediction score (**Figure 2d: Evaluation**). If a model appropriately distinguished between the labeled populations, it would assign a score near 1 to the pathogenic mutant cells and a score near 0 to the wild-type cells, and the ranked prediction score curves will be widely separated. We then selected the model with the largest separation between the labeled populations in both the training and testing wells (lack of separation in the testing wells indicated overfitting or sensitivity to batch effects) (**Figure 2d**).

Using the selected model, we chose cells for isolation from the unlabeled wells with additional control cells picked from the labeled wells. Over 1000 cells were then individually isolated into wells of several 96-well PCR plates (384 labeled, 758 unlabeled). From there, the cells underwent amplicon library construction, multiplexing, and NGS, from which their genotypes were ascertained.

With the knowledge of each cell’s genotype (n_WT_=159, n_L76P_=61, n_R94Q_=74, n_P251A_=41, n_R280H_=6), we found the total accuracy of our on-the-fly predictions to be 72.4% (50:50 =75.7%, and 10:90= 64.5%). Therefore, when training with only WT vs. a mixture of *MFN2* mutants, we could predict that individual cells were mutant from an admixture in which we were completely blind to the real genotype using only the subtle mitochondrial phenotype (**Figure 3a**). As expected, when trying to identify mutants that were the vast minority (in the 10:90 admixture), the model does correctly identify pathogenic mutant cells, but the type II error is large. **Figure 3b** shows further breakdown of all unlabeled wells by specific mutant and by whether the model correctly identified it as a pathogenic mutant (per-mutant metrics are not available in this experiment since we did not train on individual classes of mutants). The mutations most successfully recovered were R94Q and L76P, agreeing with the previous data showing those mutants as having a more severe phenotype. We recovered a similar number of false negatives of the P251A and R280H mutants, despite fewer overall numbers of each mutant, indicating that these mutants likely have a weaker phenotype—leading to a less confident prediction and exclusion from the list of cells to isolate—rather than less penetrance. To check the model quality, we evaluated the Receiver Operating Characteristic (ROC) curves and the resulting area under the curve (AUC) **Figure 3c**. The curves in red show the ROC of our model—for the cells picked from labeled (AUC = 0.94) and unlabeled (AUC = 0.74) populations.

**Figure 3.**
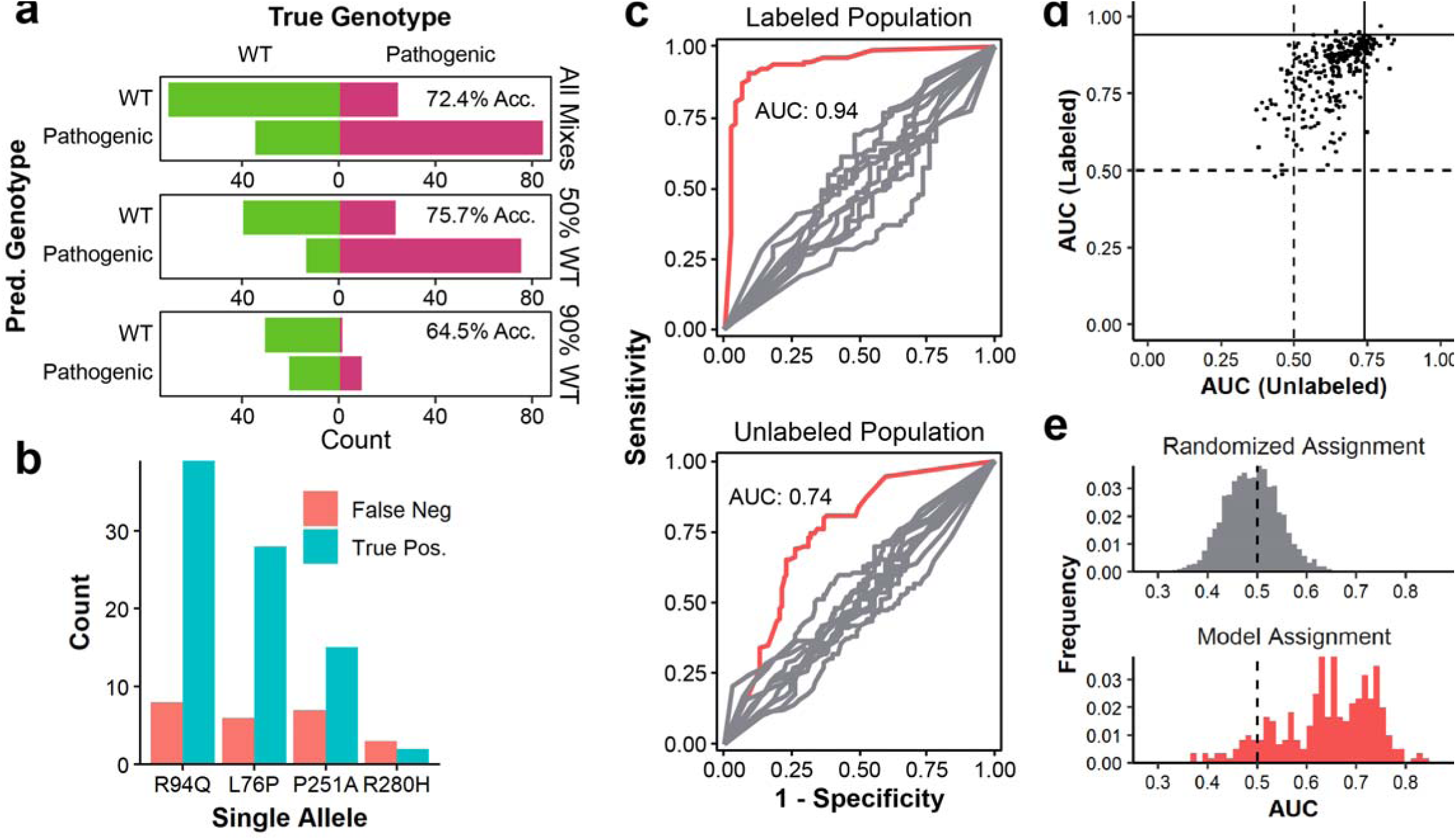
Raft-Seq Proof-of-Principle Results against MFN2 Pathogenic Mutations. **a.** Confusion bar charts of predicted genotype vs. true genotype for the main model from cells in the WT/Pathogenic mixture. The top bar chart represents all cells isolated from the mixture, while the bottom two are a breakdown of the first according to whether the cells were isolated from the well containing 50% or 90% wild-type cells. **b.** A bar chart of picked mutants separated by allele and counting the final status of individual cells. **c.** ROC Curves generated using the best model identified *a posteriori* separated by data generated from cells picked from control pure wells (upper) and from mix wells (lower). The red curves are the experimental results and the grey curves are a control generated by random shuffling of labels. **d.** A scatterplot showing the performance of all 290 models in detecting mutants in a mixture of wild type cells and four pathogenic mutants. Each point represents a single model and its position is determined by its ability to distinguish cells in the labeled control wells and cells in the mixed wells. The vertical and horizontal lines represent the AUCs of the model that was used to choose cells for isolation. **e.** A histogram of AUCs for models detecting mutants in a mixture of wild-type cells and four pathogenic mutants. On top, a histogram of AUCs generated from randomly assigning models is shown as a comparison.

We also took a larger view and assessed the quality of *all* models generated, not just the one used to select cells for isolation. **Figure 3d** shows a scatterplot of AUCs (labeled populations vs. unlabeled populations) and **Figure 3e** shows the AUC distribution of all models applied to unlabeled populations. Importantly, the majority of models had discriminatory ability. This entire experiment was repeated successfully using a mixture of all six mutants, and histograms and scatterplots of the models generated are shown in **Figure S3.**

### Identification of Weak Pathogenic *MFN2 Mutants*

Our pipeline also has the ability to recognize relatively weak phenotypes. As shown earlier, of the six pathogenic mutants, the R280H and P251A appear most like benign mutants and wild type (**Figure 2c**). Despite the subtle phenotype, we recovered R280H cells in the experiment above (**Figure 3b**), though comparably fewer than cells with other mutations. Given those results, we explored the ability of Raft-Seq to isolate cells with the R280H mutation compared against *MFN2* wild type. **Figure 4a** shows images of R280H mutation-harboring cells in the microraft plate. As a baseline measurement against more conventional methods of cell separation, we ran separate samples of a *MFN2* WT and R280H mutant cell lines each stained with MitoTracker through flow cytometry (**Figure 4b**). While there is variation between the two in the MitoTracker intensity, there is too much overlap to separate a mixture (**Figure S4** AUC = 0.60).

**Figure 4.**
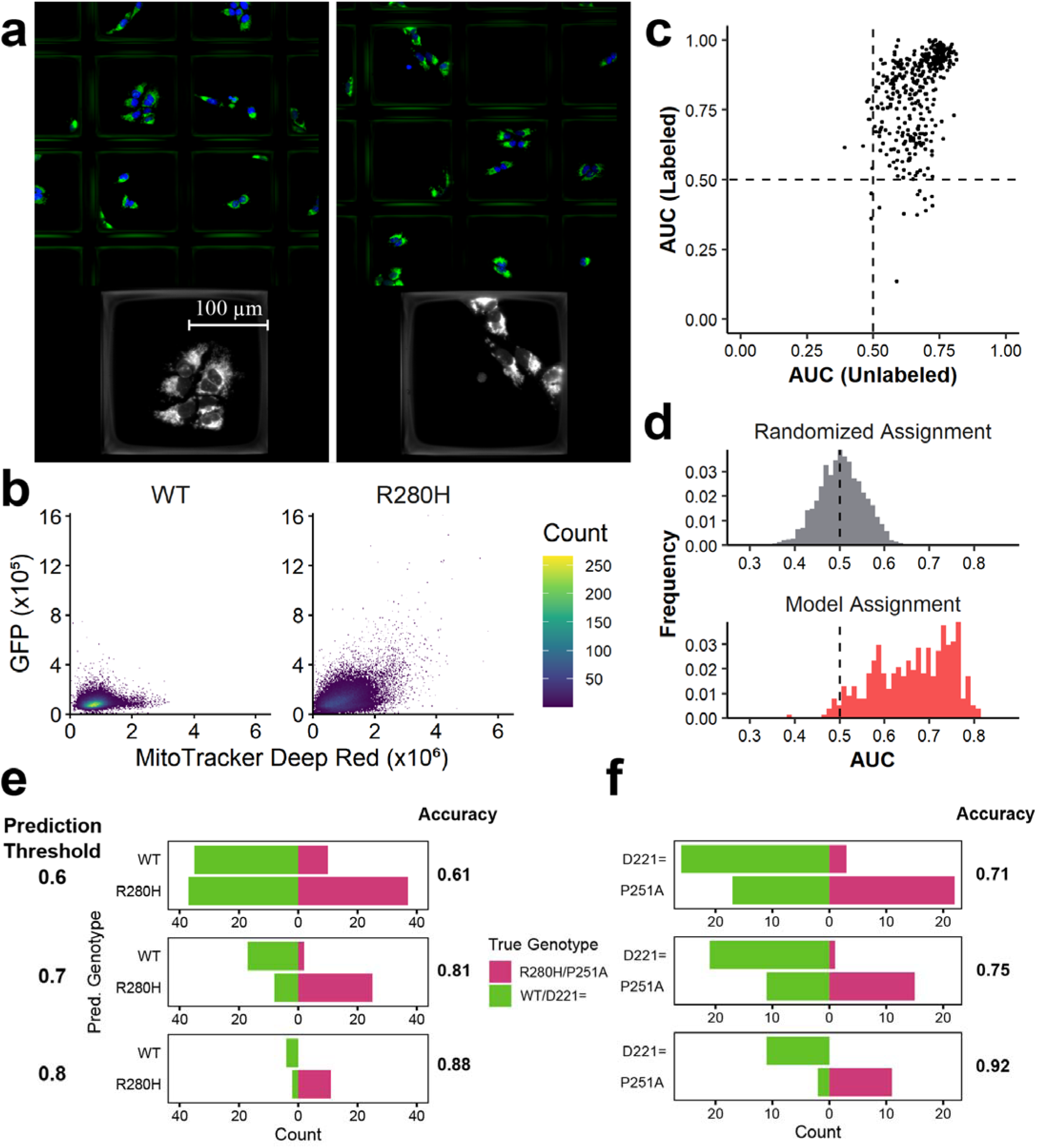
Raft-Seq can Predict *MFN2* WT and *MFN2* R280H mutant cells which have nearly undetectable Phenotypic Differences. **a.** Images of the R280H mutant and *MFN2* WT cell line in the microraft plate. **b.** Side-by-side flow cytometry scatter plots of the *MFN2* WT and R280H mutant cell (GFP was present equally in both cell lines). **c.** Scatterplot showing the performance of all 433 models in detecting mutants in a mixture of wild type cells and the R280H mutant. **d.** A histogram of AUCs for models detecting mutants in a mixture of wild-type cells and the R280H mutant. On top, a histogram of AUCs generated from randomly assigned models for comparison. **e,f.** Accuracies and confusion bar charts of the predictions from the un-labeled wells, when only picking cells with prediction scores ≥0.9 (≤0.1), ≥0.8 (≤0.2), and ≥0.7 (≤0.3). **e.** uses the model to predict R280H against WT *MFN2*, while **f.** uses the model to predict P251A against D221=.

We then followed the Raft-Seq process described above, replacing the mixture of *MFN2* mutants with just R280H mutant cells, and therefore training the models directly on the weaker phenotype. Additionally, the experiment was done with different culture densities and cross-compared to confirm that culture conditions had no ‘residual phenotype’ that impacted the models prediction. **Figures 4c** and **4d** show histogram and scatterplots for the resulting models in this experiment and that Raft-Seq discriminated between the R280H mutant and the *MFN2* wild type (AUC of picked model 0.72, best models > 0.8, n_WT_ = 265, n_R280H_ = 205). We also ran a similar experiment comparing the P251A pathogenic mutant cell line—the other pathogenic mutation with a weaker phenotype—to the D221= benign mutant cell line. Cells containing the synonymous substitution D221= act as another control for point mutations instead of using WT. **Figure S5** shows that the resulting model, when applied to a mixture of the cell lines, was able to discriminate between the two (AUC = 0.8, n_D221=_ = 103, n_P251A_ = 65). The AUCs mentioned above apply to all the unlabeled cells that were physically picked. If we limit our results to cells whose class the model is increasingly confident about, then the accuracy of the model gets increasingly better (**Figure 4e, f**). For cells that the model is at least 80% confident about (prediction score ≥ 0.8 or ≤ 0.2), we get high AUCs and accuracies in both the R280H/WT and P251A/D221= experiments (AUC = 0.94,0.98 Acc = 0.88,0.92 n=17,24).

We have shown that Raft-Seq can accurately predict genotypes from strong mitochondrial point mutants as well as weak mutants in *MFN2* and predict them as part of a mixed culture where there was no *a priori* knowledge of individual cell’s genotypes. The model’s predictions were realistically tested by isolating single cells and genotyping them to reveal the method’s accuracy.

### Modeling Analysis

Multi-feature models were necessary for accurate predictions, warranting exploration into the importance of individual features and the performance impact associated with them. Using data from the experiment containing wild-type *MFN2* cDNA and four pathogenic mutants, we generated additional models by varying the number of features used, sampling from the 8 features in the pick model (**Table S1`**). Histograms showing the distributions of the AUCs, separated by the number of features are shown in **Figure 5a.** When more features are added, the models can better distinguish between the populations of cells. We also examined the importance of individual features in building models (**Figure 5b-d**). We used the original feature data (**Figure 1**) and generated Kruskal-Wallis □^2^ for each feature between the wild type and pathogenic mutant cells from labeled populations. A ranking of the 16 features associated with the highest □^2^ are shown in **Figure 5b**. We compared the resulting AUCs from models built off each specific feature alone (**Figure 5c**), and to the resulting AUCs from models built from the 16 features minus one (**Figure 5d**). Importantly, we find that single features *are not* the key to discriminating these clinically relevant pathogenic *MFN2* point mutants. Instead, small numbers of relevant features can inform a useful model.

**Figure 5.**
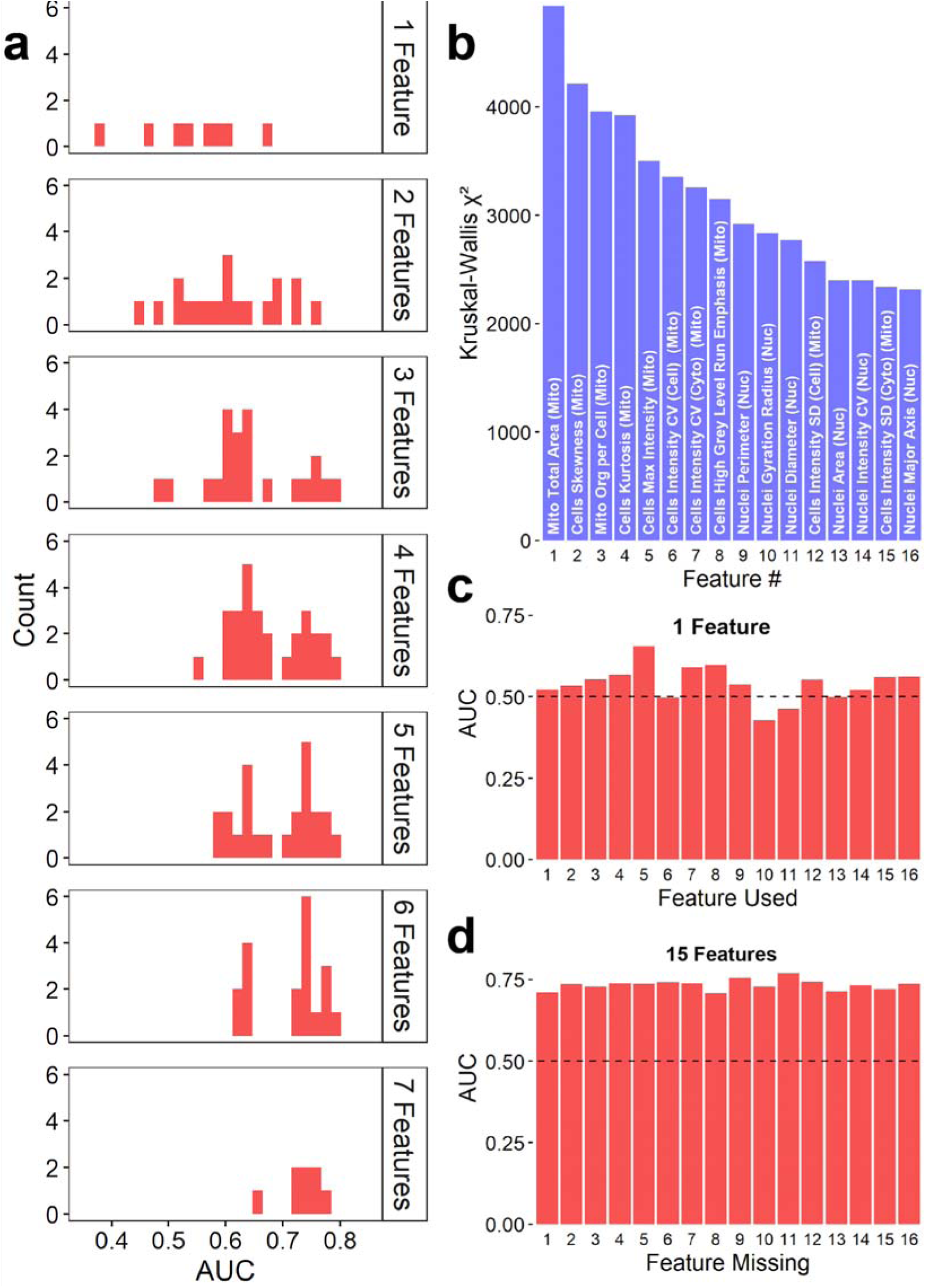
*MFN2* Genotype Prediction does not depend on specific features. **a.** Histograms of AUCs resulting from models built from combinations of a set of 8 features used when picking, arranged in panels split by the number of features used in each subset. **b.** Bar chart of the Kruskall-Wallis □^2^ values for the 16 features with the highest values. “Mito” and “Nuc” indicate features measured from mitochondrial and nuclear stains, respectively. **c.** Bar chart of the AUCs resulting from models built out of each individual feature. Note that the most significant feature was unable to produce a good model alone. **d.** Bar charts of the AUCs resulting from models built using all but one feature (leaving 15 features). All AUCs listed indicate the performance of the model trained from labeled data on their ability to predict *unlabeled* cells in admixed conditions.

### Scanning Mutagenesis with *MFN2* VUS

While the previous experiments were done by over-expressing a mutant *MFN2* cDNA, we also sought to study editing of an endogenous gene. As a proof of concept for endogenous mutations, we first used an existing U2OS line with mutations in the mitochondrial primase *PRIMPOL* ^42^ and found that it alters mitochondrial morphology. We then followed the Raft-Seq process described above and verified that the platform performs well for an endogenous genetic perturbation (AUC 0.90, data not shown). We next tested Raft-Seq against a series of (mostly VUS) mutations in *MFN2*. We constructed a CRISPR-Cas9 gRNA library targeting different exonic regions of *MFN2* (CRISPR tiling ^43^) on sites near known ClinVar variants (**Figure 6a**). The library was delivered by lentivirus to U2OS cells with a dox-inducible Cas9 construct, and the cell population was split in two, with only half receiving doxycycline. DNA samples were collected at intervals over a ten-day period for survival screening, and a DrugZ analysis ^44^ revealed no significant representation differences (**Figure S6**)—so while mitochondrial changes are observed, those don’t result in large viability shifts.

**Figure 6.**
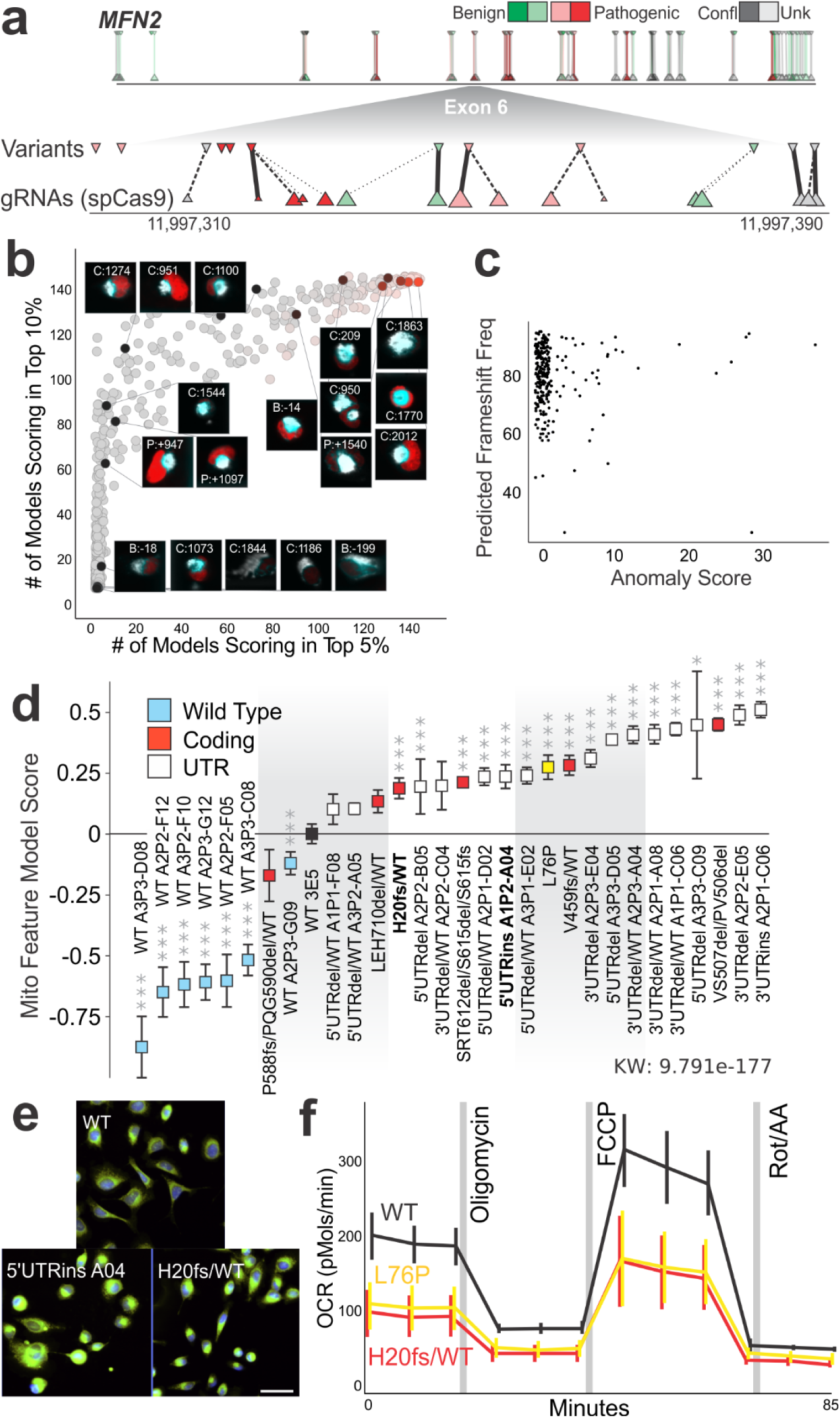
Endogenous Scanning Mutations in *MFN2* simulate Variants of Uncertain Significance. **a.** Diagram of coding sequence with ClinVar variants (downward facing triangles) and their closest gRNA cut sites (upward facing triangles). The upper line shows the entire genomic context of the MFN2 gene, while the lower inset magnifies the variants and gRNA cut sites for exon 6. Lines indicate the ‘partner’ gRNAs for each ClinVar mutation, where thicker lines indicate closest association. **b.** Scatterplot of genotyped cells. x and y axes are the number of models placing that cell in the top 5% or 10% of all cells, respectively. A sample of cells are labeled with the fluorescent image taken during the screen and the gRNA present within the cell (labeled by the nucleotide position before the start codon [B], within the coding region [C], or after the stop codon [P], also see **Table S5`**). **c.** Scatterplot of predicted frameshift frequency vs. mitochondrial anomaly score for all the gRNAs recovered from the screen**. d.** Established isogenic lines of U2OS cells with the specific *MFN2* variant(s) rank ordered based on their phenotypes. More “WT-like” phenotypes (likely benign) are shown to the left while more “mutant-like” phenotypes (likely pathogenic) are shown on the right. Bars show average with 95% confidence intervals (n=18-32 well/plate replicates, L76P mutant is shown in yellow and the parental control clone WT 3E5 is shown in black). **e**. Example images of isogenic U2OS cells with either a control gRNA that didn’t cut, 5’UTRins A04 isogenic line, or H20fs/WT isogenic line. The frameshift-mutant (H20fs/WT) has similar mitochondrial aggregation to the L76P mutant (Figure 1), while the UTR mutant has a distinct but subtle mitochondrial morphology. **f**. Metabolic analysis of OCR (oxygen consumption rate) for WT, L76P, and the H20fs/WT lines. OCR is measured in four phases, first basal, then mitochondrial toxins Oligomycin (ATP synthase inhibitor), FCCP (uncoupler), and Rotenone (electron transport inhibitor).

Then in a Raft-Seq screening experiment, 40,000 cells were imaged, and mitochondrial anomaly detection models were trained on the feature data of -Dox (no Cas9) cells and deployed to the data of +Dox (Cas9 induced) cells to infer which cells had abnormal mitochondrial phenotypes. Unlike previous modeling techniques, training data of known mutants was not included. The most highly anomalous cells had some feature similarity to the strong pathogenic mutants (**Figure S7**). We selected 1,659 cells for isolation, which we then genotyped to identify the gRNA for each cell (**Figure 6b**). The result is a rich dataset where each of the single cells are measured for phenotypic features (from confocal imaging) *and* the corresponding gRNA. Since individual gRNAs have different efficiency, we presumed that frameshift edits to *MFN2* would be more likely to result in the strongest phenotypes. We compared each cell’s mitochondrial anomaly score to the probability of the gRNA inducing a frameshift mutation (calculated with Indelphi ^45^). We find that nearly all gRNAs with a high anomaly score also had a high predicted frameshift mutation rate (**Figure 6c,** p-Value at 3 frameshift thresholds: 90% = 0.0013 [n 42], 80% = 0.05 [n 111], 70% = 0.37 [n 39]).

Next, we harnessed the capabilities of Raft-Seq to analyze a library of perturbed cell lines. We took the same population of *MFN2* gRNA library-infected cells and chose cells for isolation based on individually identified mitochondrial morphology using a parallel pipeline. We captured the cells intact and alive into tissue culture plates and grew them clonally as isogenic lines. This allowed enough genomic material to do two rounds of sequencing—first to find the gRNA present in each cell and second to examine the gRNA target site—and use the cells for downstream analyses. Most of the isogenic lines with Cas9 edits were in the UTRs, while 6 affected the protein (**Table S2`**). For example, the isogenic H20fs/WT had a gRNA targeted to the first translated exon, which resulted in 7bp deletion then a frameshift. We analyzed the isogenic cells (arrayed) to quantitatively assess the level of mitochondrial abnormality (**Figure 6e**). Three dozen of the individual mitochondrial or TMRM features (**Table S3`**), as well as all multi-feature models, showed significant differences between the set of WT clones and the mutated clones (**Figure S8**).

We performed additional validation experiments on two of the clones, one UTR-mutant and one coding-mutant (**Figure 6f**). We found that these isogenic endogenous mutations were separatable from a WT clone in our Raft-Seq mix assay (AUC = 0.90, **Figure S9**). We also analyzed the set of *MFN2* isogenic lines using the Seahorse XF assay, which analyzes key metabolic processes reflective of mitochondrial health (**Table S4`**). Both the H20fs/WT and the L76P mutant had reduced basal oxygen consumption rate compared to WT, indicating a mitochondrial deficit from the mutations (**Figure 6g**). Overall, we showed pooled cellular screening with Raft-Seq and its ability to generate isogenic lines with mutant phenotypes, including novel mutations that have not previously been studied.

## Discussion

We have developed a method, Raft-Seq, to efficiently screen many genetic variants based on their impact on a cell’s phenotype. We have shown that it can effectively discriminate between wild-type cells and cells containing different pathogenic point mutations of the *MFN2* and *PRIMPOL* genes. Though we utilize a particular individual model for the selection of cells for isolation, we found that most of the models that we generate can identify pathogenic mutant cells. Since our feature selection process is only mildly dictated by the actual phenotype that we are looking for, our ability to predict a cell’s genotype comes from quickly generating complex computational models. This will enable the pipeline to exploit many more phenotypes that the scientific literature is not yet familiar with, therefore making almost any gene in the genome amenable to this functional screening and thus enable the re-cataloging of VUS as benign or pathogenic.

We developed Raft-Seq primarily as a screening platform to work in concert with Deep Mutational Scanning ^46^, a method for creating a library of every single possible mutation in a gene. Our results show that such a screen would correctly call pathogenic variants, since we were able to recover pathogenic *MFN2* mutants from a mixture with wild-type cells. However, our results also show some limitations. For one, based on the accuracy difference between the two unlabeled wells which were at different proportions, our ability to recover pathogenic variants appears to decrease the smaller proportion of pathogenic variants present. This would indicate that such a screen would perform significantly better on genes for which mutations are more likely to result in a pathogenic variant, but this could be countered with a higher n. Second, we were able to identify several mutants (L76P, R94Q)—which had more extreme phenotypes—more frequently than other mutants, meaning that a scaled-up screen would most likely result similarly, and more extreme variants would be overrepresented. However, because we were able to identify the weaker mutants (R280H, P251A) with relative ease when they were not mixed with any other mutants, we can assume that overrepresentation of specific variants is not due to the absolute strength of the resulting phenotype. Instead, it is due to the relative strength when compared to other variants. Penetrance may also play a role since weaker phenotypes may have a strong phenotype in some cells but have low penetrance in the population.

We were also able to see how Raft-Seq would perform in the discovery of novel variants and morphology. Our anomaly detection models were able to identify similar mitochondrial features to the pathogenic variant cells, and this anomalous morphology correlated with a high predicted frameshift frequency from the gRNA contained in the cell. Lastly, we used Raft-Seq to generate isogenic cell lines directly from the primary screen. We analyzed these clonal/isogenic lines to find consistent mitochondrial phenotypes and blunted metabolic responses. This process presents a large gain over other phenotyping screens, since we can generate cell lines that can be fully genotyped and assayed further, while being able to track that line to the specific image and features from the original screen.

Since Raft-Seq isolates cells for sequencing individually rather than in pools, we have a few advantages over other platforms. For one, we find the specific genotype of every isolated cell, rather than perform batch measurements, meaning that we can find effects of combinations of perturbations. We can also theoretically have as many categories in our machine learning models as cells, though using more than a handful of features causes the modeling and analysis to deteriorate, likely due to overfitting (but is countered by higher n).

An advantage of this approach is the flexibility that comes from using machine learning to identify phenotypes. Since we can easily combine features for more accuracy, we can theoretically screen for any visible phenotype, given a strong enough signal provided by staining or other fluorescence. Unsupervised learning methods, like clustering, are also possible and allow for a simpler setup since no labeled wells would be needed.

Since the raft identifies the location of the cell stably over time, we can do on-the-fly training from the entire experiment, then go back and pick those cells. We have also found that Raft-Seq can be used to apply past training (labeled) data to future experiments. A normalization schema allows for accretion of previous training data to utilize it in future models (allowing a gain of accuracy in identifying specific phenotypes over time). Anomaly detection models need previous data, to capture as much “normal” cell data as possible.

Raft-Seq is slower than flow cell sorting, possibly limiting scalability. All the image-based techniques require time for imaging, and this system is no exception. As evidenced by the recent genome-wide screen ^25^, the throughput of this system is comparable, and the number of cells captured by that system is slightly lower over an experiment of comparable scale. Regardless, even higher throughput is still advantageous. There are fewer limits in terms of the number of cells able to be imaged, so by increasing the pool of isolation candidates, cells are selected with a higher level of confidence. There are two steps in Raft-Seq that are done manually but will soon be automated. The first is image quality control which can be automated by a convolutional neural network, similar to the previous microraft experiments ^33^. The second is model selection, which can be automated by choosing among several model performance metrics on the labeled data.

We are optimistic that with increased scale, Raft-Seq can be used to sensitively find variants across a wide breadth of perturbed cellular phenotypes. An increase in scale would provide more training data to make models more accurate and identify more cells to which the models assign a high prediction score, which we have shown to raise overall accuracy. Altogether, we have shown that using various cell lines (HEK293, U2OS, A549), with multiple types of genetic perturbation (endogenous disruption with Cas9, overexpression of mutant genes), we can deploy flexible machine learning (logistic regression, decision trees, SVM, neural networks) from data within an experiment (on-the-fly) or from previous experiments (pre-trained) to select clinically relevant point mutations in a screening setting. Since this system captures the cells individually, we can carefully evaluate many ‘selection criteria’ by generating a variety of different machine-learning models and finding how well they perform. This technique will be an important tool in the advancement of precision medicine to identify variants relevant to disease.

## Methods

All custom software can be found at https://gitlab.com/buchserlab/FIVTools

### Cell Culture and Transfection

Human osteosarcoma (U2OS, ATCC HTB-96) cell lines were maintained in McCoy’s 5A Modified Medium (16600082, Gibco, Gaithersburg, MD, USA) supplemented with 10% fetal bovine serum (FBS) (16000044, Gibco). Human embryonic kidney (HEK) 293T cells (CRL-11268, ATCC) were cultured in Dulbecco’s Modified Eagle’s Medium (11965-092, Gibco, Gaithersburg, MD, USA) supplemented with 10% FBS (16000044, Gibco, Gaithersburg, MD, USA), 1% Penicillin-Streptomycin (15140122, Gibco) and 1% non-essential amino acids (11140050, Gibco).

All cell lines were maintained in T75 tissue culture flasks in an incubator at 37°C, 5% CO_2_ and they were observed daily for growth and overall health. Once confluent, cells were passaged using 0.25% Trypsin-EDTA 1x (25200056 Gibco, Gaithersburg, MD, USA) at a sub-cultivation ratio of 1:10. Live cell counting was performed with the BioRad TC20 automated cell counter. Centrifugation of cell cultures was performed at 1200 rpm for 3 minutes. Lentiviral infection was performed in T75 flasks when cells were 85% confluent. STR profiling, to confirm cell type, was performed using NGS-based analysis by the Genome Engineering and iPSC Center (GEiC) at Washington University in St. Louis. Testing for mycoplasma was performed bi-annually. For all experiments in this paper, either 100×100 or 200×200 micron quad reservoir plates containing 48,000 (12,000 cells per quad) and 36,000 cells (9,000 cells per quad), respectively were used. Prior to plating, microraft plates were prepared by rinsing with 1mL PBS 3 times with 3-minute incubation periods. Cells were added in 200µl media to aid in distribution, then plated and incubated overnight (14-16 hours).

### Virus Production and *MFN2* Single Mutant Line Creation

*MFN2* lentiviral expression plasmids were cloned into the CCIV lentiviral plasmid with a GFP marker ^39^. In preparation for lentiviral packaging, 8.0 x 10^5^ HEK293T cells were plated into each well of a six well plate and incubated at 37°C overnight. The cells were then transfected with TransIT Lenti-transfection reagent (MIR 6600, Mirus Bio, Madison, WI, USA) using an envelope plasmid (pVSVg: Addgene plasmid # 8454), a packaging plasmid (psPAX2: Addgene plasmid # 12260), and each individual MFN2 expression plasmid in a mass ratio of 0.5/1/0.5 respectively for a total of 2µg. After 48 hours, media was collected, centrifuged, and sterile filtered before being concentrated (Lenti-X Concentrator 631232 Takara Bio, Kusatsu, Shiga, Japan). Concentrated virus was resuspended in 200µL 1xPBS per well, collected, and stored at -80°C.

To create stable MFN2-mutant expressing lines, T75 flasks containing 6 million U2OS cells were infected with 70µL of concentrated lentivirus at an MOI>1 and polybrene was added (NC9840454 Santa Cruz Biotechnology, Texas) at a final concentration of 10µg/mL. They were then incubated for 24 hours, after which the virus-containing media was removed and replaced with fresh, virus-free media. Cells were taken to the Washington University Siteman Flow Core for fluorescent sorting on the Sony Synergy HAPS1, 100-micron sorter. Cells were sorted based on viability and GFP expression (since no puromycin selection was performed, the fluorescent signal from the GFP in the MFN2 plasmid was used to determine transgene expression). GFP expression levels were compared within and across generated cell lines to ensure population purity and comparable fluorescent expression levels. The PRIMPOL KO U2OS cell line was received from the Vindigni lab and was produced by the GEiC.

### CRISPR/Cas9 gRNA Library Infection and Induction

A dox-inducible Cas9 (iCas9) U2OS cell line was generated via CRISPR-mediated homology directed repair. The Cas9 protein, gRNA, and donor construct were introduced via nucleofection. Isogenic iCas9 clones were isolated using the Cell MicroSystems CellRaft Air System and then propagated for further experiments. Presence of the construct was validated via junction PCR ^47^ prior to propagation. Puromycin-resistant *MFN2* scanning gRNA libraries were generated and cloned by the Washington University GEiC. Lentivirus was produced (see Virus Production above) and used to infect iCas9 U2OS cells at an MOI of <0.2 followed by 8µg/mL puromycin selection for seven days. The cells were then allowed to grow in fresh media. At 60-70% confluency, Doxycycline (Cat#: D9891-1G, Millipore Sigma) was added at a final concentration of 2µg/mL. The cells were incubated at 37°C for 48-60 hours before proceeding with staining and imaging.

### Staining and Microscopy

The following vital dyes were used; DNA labeling/nuclei (Hoechst, Thermo Fisher H3570), mitochondria (MitoTracker Deep Red, Thermo Fisher M2246), and mitochondrial membrane potential (Tetramethyl Rhodamine methyl ester TMRM, Thermo Fisher I34361). MitoTracker and TMRM were incubated for 40 minutes at concentrations of 0.5 and 1µM respectively. Hoechst was incubated for 15 minutes at a concentration of 10µg/mL (16.2µM). Each plate was rinsed twice with culture media prior to imaging. Images were captured using a 20x 0.45 NA objective in the Cytiva INCell 6500HS Confocal microscope. Exposure times for Hoechst (405 nm) and TMRM (561 nm) averaged 0.15 seconds while MitoTracker Deep Red (642 nm) averaged 0.05 seconds. Confocality was used in the 405 and 642 wavelengths to decrease the background fluorescence of the CytoSort raft plate. Each field-of-view overlapped by 12% of their area. Imaging settings were held constant throughout the course of an experiment. Following imaging, an extra 500µL of cell culture media was added to the CytoSort raft plate (additional liquid helps the Cell MicroSystems Air System isolate microrafts).

### Image Analysis and Quality Control

Image tracing and feature extraction was performed using Cytiva’s INCarta software. Mitochondrial puncta were identified (within 20µm of the nuclei using the ‘networks’ algorithm) and quantified for each cell as were a set of texture features. Raft coordinates were recorded for each cell (using FIVTools/ CalCheck, included in the GitLab repository). Images were also curated semi-manually (via FIVTools/ CalCheck) to ensure that out-of-focus images were excluded. The cell feature dataset was joined with the image quality data and raft position mapping data described above by custom software (via FIVTools/ main window). Post tracing quality control was performed with each dataset in Tibco Spotfire Analyst. First, aberrant tracing artifacts were excluded based on nuclear area, nuclear form factor, and proximity to the raft’s edge. Next, non-nuclear debris and dead nuclei were excluded by gating on nuclear area, intensity, and cell intensity. Rafts with too many cells (>6) or a fiduciary marker were excluded. This filtered set of cells was used as the input for machine learning downstream.

### Machine Learning and Model Generation

After exporting the quality-controlled cell-based feature table, we built a machine learning model that could distinguish pure populations of WT cells from pathogenic mutant cells. The models were trained real-time on the day of the experiment since we were working with live cells which needed to be physically picked within the next few hours to maximize viability. A variety of machine learning platforms (Microsoft AzureML Studio, Tibco Spotfire, Tibco Statistica, and H2O.ai.) and algorithms were employed to predict an unknown cell’s genotype class. Generally, logistic regression was performed in Spotfire and random forests, boosted trees, support vector machines, gradient boosted machines, and artificial neural networks were trained in the other platforms. Models were evaluated on labeled populations that were withheld during training. Based upon model performance on the testing dataset, a model was selected and deployed to the unlabeled cell populations. Starting with the strongest prediction scores, a list of cells with raft locations was generated.

### Cell Capture and DNA Extraction

Cells were isolated using the Cell MicroSystems CellRaft Air System. CytoSort raft plates were received from Cell Microsystems (Durham, North Carolina). Given a list of raft coordinates, the Air System used a needle to eject each individual raft and transfer the raft to a semi skirted 96-well PCR plate (1402-9200, USA Scientific) via a magnetic wand. Each well of the PCR plate contained 5µL extraction buffer (molecular grade water with 10mM Tris-HCl (pH 8.0), 2mM EDTA, 200 µg/mL Proteinase K, and 0.2% TritonX-100). Raft isolation was confirmed twice through post ejection imaging of the raft location and through visual inspection using a Leica S8AP0 dissection scope. Genomic DNA was extracted in a thermocycler immediately following raft isolation by incubating at 65°C for 15 minutes then 95°C for 5 minutes.

### Single-Cell DNA Amplification

Amplification of single-cell DNA prior to library preparation consists of two separate amplifications. An initial preamplification is conducted using extracted DNA with KOD Hot Start DNA Polymerase (71842-4, Millipore Sigma, Burlington, MA, USA) according to manufacturer’s instructions using all 5µL of extracted DNA in a total reaction volume of 20µL. Pre-amplified product was processed through an AMPure XP (Catalog: A63882, Beckman Coulter, Brea, CA, USA) bead clean up according to the manufacturer’s instructions using 10mM Tris-HCl pH 8.5 as elution buffer. The second amplification uses the cleaned template and BioLine MyTaq HS Red Mix 2x (C755G97,Meridian Life Sciences, Memphis, TN, USA), according to manufacturer’s instructions, including 5% by volume DMSO. Primers in the second amplification contained universal 5’ tags to be compatible with Illumina library preparation (Forward tag: 5′- CACTCTTTCCCTACACGACGCTCTTCCGATCT-3′, Reverse tag: 5′- GTGACTGGAGTTCAGACGTGTGCTCTTCCGATCT-3′).

For amplification of *MFN2* cDNA, primers amplifying the entire cDNA were used in the first amplification step, followed by multiplexed amplification of two specific regions containing the relevant mutations. Genotyping of the RFP-GFP cells used multiplexed primers that amplified specific regions in both the RFP and GFP regions. All primers are listed below in table 2.

**Table 2.**
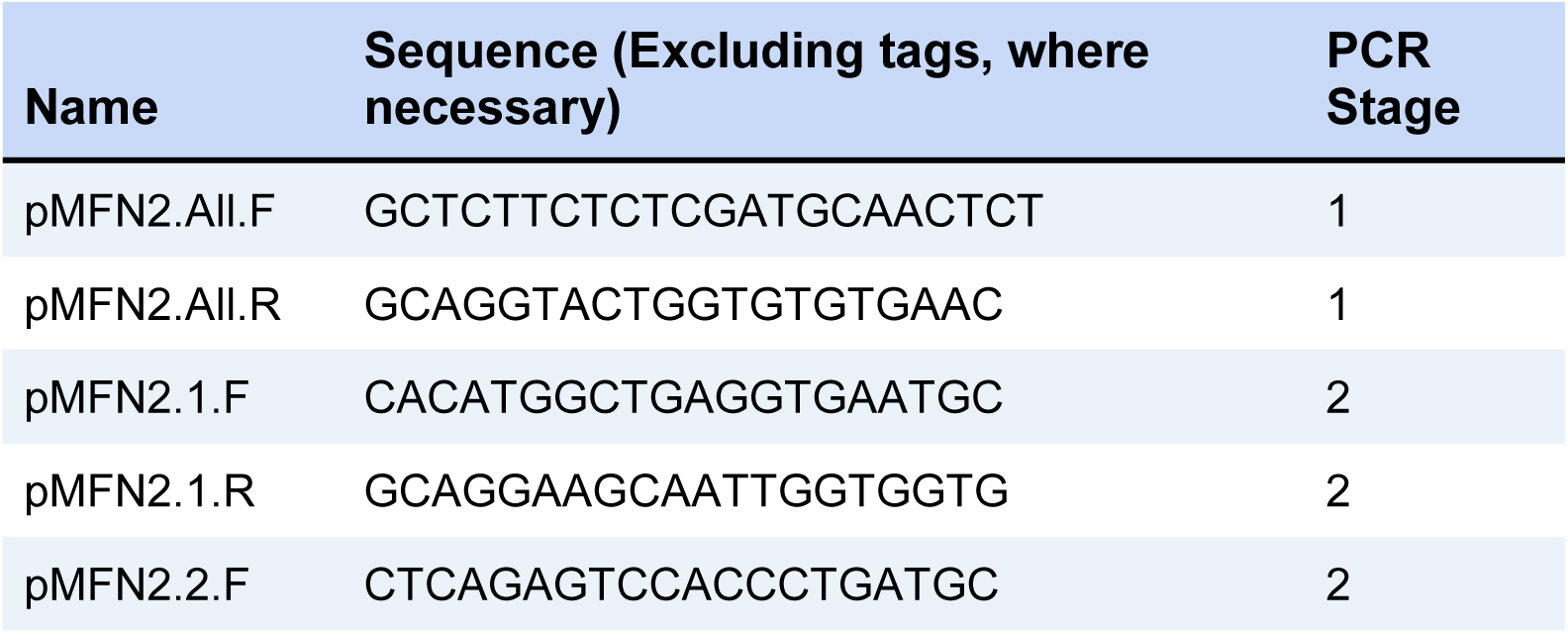
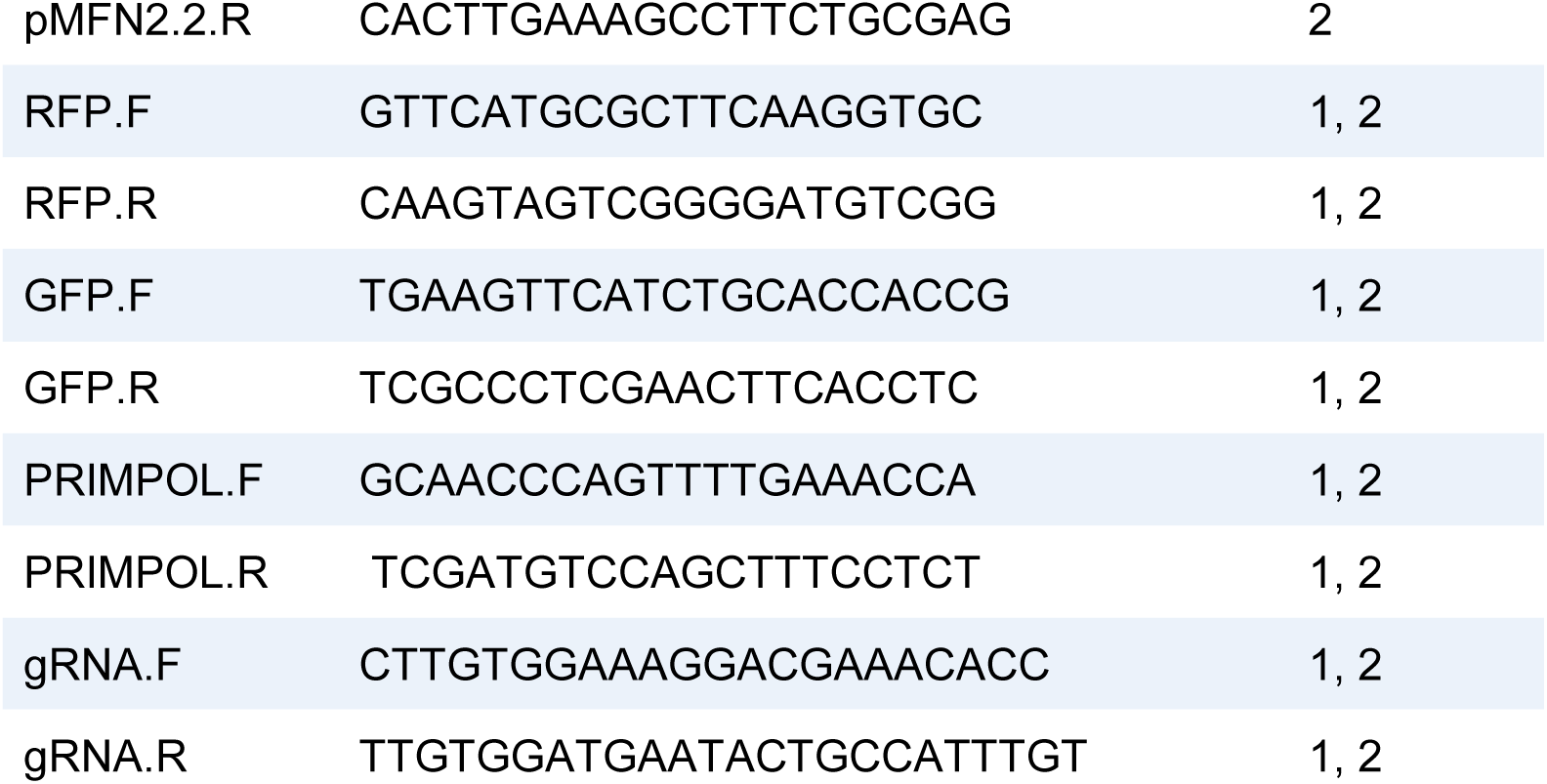
Genotyping Primers.

### Illumina Library Preparation

These methods are expanded from Connelly et al. and Bell et al. After amplification with universal primers, each plate was amplified with specific forward and reverse Illumina index primers that indicate the PCR plate position and a unique plate ID. PCR amplification was performed with BioLine MyTaq HS Red Mix 2x (C755G97,Meridian Life Sciences, Memphis, TN, USA) according to the manufacturer’s protocol, pooled, and then cleaned using AMPure XP bead (A63882, Beckman Coulter Life Sciences, Indianapolis, IN,USA) cleanup procedure in the original amplification. DNA was quantitated on a NanoDrop One Spectrophotometer (Thermo Scientific, ND-ONE-W) before being submitted to the Center for Genome Sciences and Systems Biology (Washington University) to generate 2×250 reads on the Illumina MiSeq platform.

### Sequencing Analysis

Illumina paired reads were demultiplexed by the core facility and FastQ files were returned. The rest of the analysis was performed with laboratory software available on Gitlab (FIVTools/ LA, “Library Aligner”). Reads were joined and trimmed, then aligned with small sequence fragments at the genetic sites of interest containing the sequence to mutant or WT alleles. The result was a ‘counts’ table that gave the number of reads containing each 20-mer for each well. 20-mer search fragments are listed below in **Table 3**. After accounting for isolation and genomic amplification errors, around 80% of the isolated cells genotypes were captured.

**Table 3.**
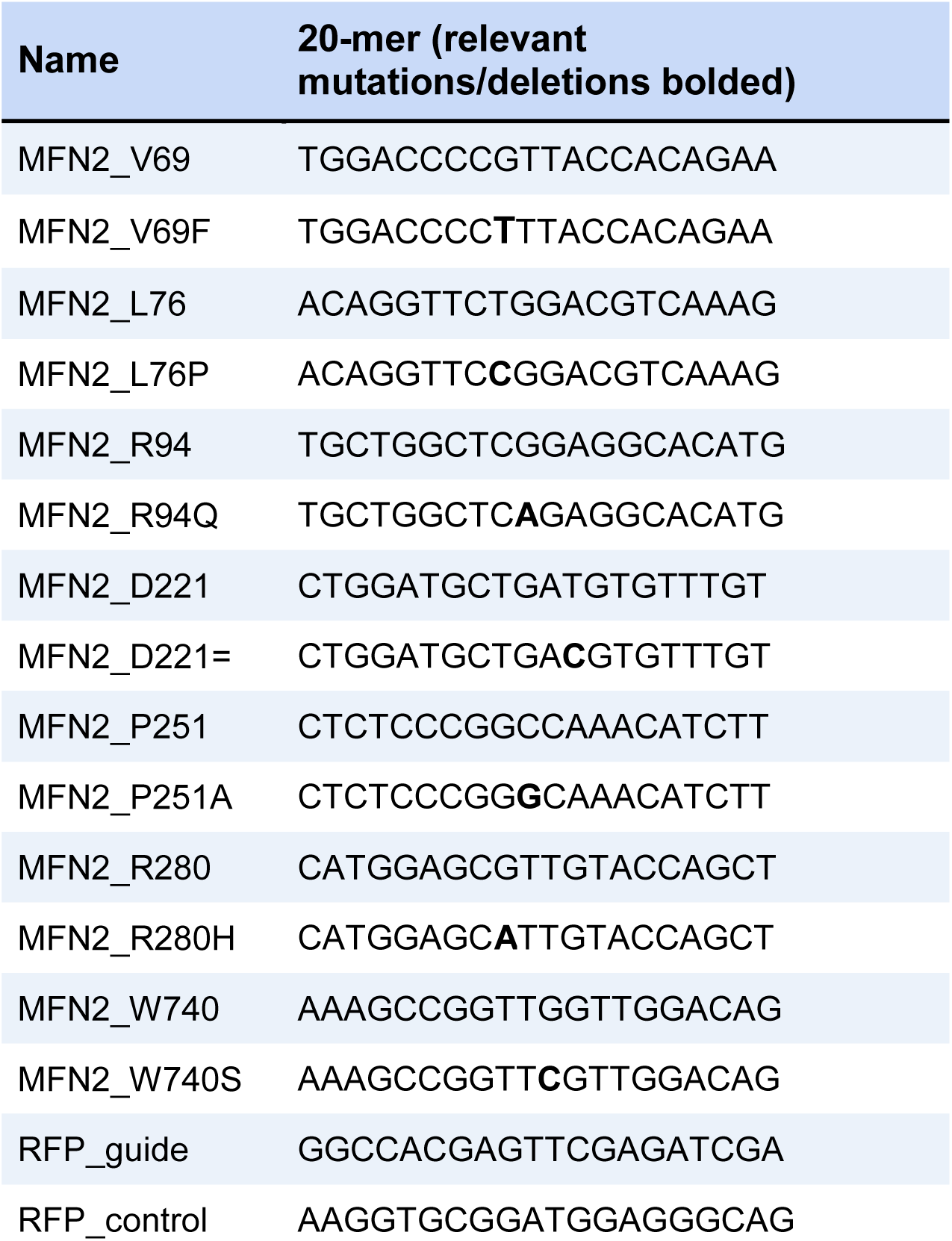
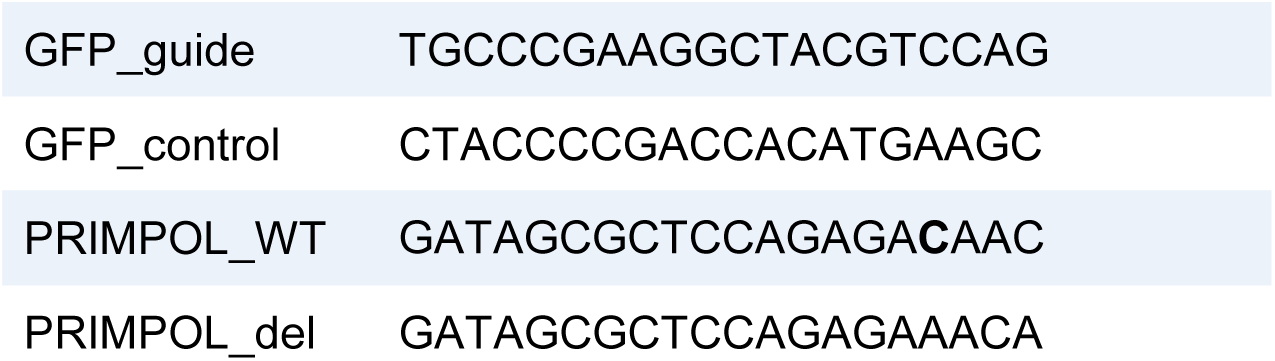
20-mer fragments.

For *MFN2* cDNA genotyping, each mutation locus was given a %mutant score calculated as the number of mutant reads divided by total number of reads at that locus. Cells were designated as wildtype if no locus had >50% mutant score, otherwise they were designated as a specific mutant based on which locus had the highest mutant score (ambiguous cells were excluded). Lastly, a flat file was exported containing each picked raft and its assigned genotype.

Using our custom software (FIVTools/ AUC), we joined the modeling and genotyping flat files to find overall accuracy and generate ROC curves for each model. We also generated ‘noise’ ROC curves by shuffling the assigned genotypes. Prediction scores between 0.4 and 0.6 were filtered out (scores near 0.5 meant the specific model was unable to classify these cells). For the data presented in **Figure 4e,d**, this threshold was further adjusted.

### Isogenic Line Production

For clonal cell growth, single live cells were isolated by the Cell Microsystems Air System into 96-well tissue culture plates (TPP 92096), containing 200µl of media per well. As the isogenic lines grew, the entirety of each well was passaged into a plate of larger size (96 to 24 to 12 to 6 well plates from TPP) after reaching ∼50% confluency. It took 2 weeks to go from single-cell to 50% confluency in the 96-well plate, and during that time wells were checked for contamination and media level every 2 days. After the cells were plated in the 6-well plate, one third of the cell suspension was taken through DNA extraction for genotyping. One third was frozen down for long term storage, and the remaining third was kept for downstream experiments. The entire process took ∼2 months to go from single cells to frozen stocks/genotyping data. Genomic DNA samples were initially genotyped to determine the gRNA(s) present (as described in the preceding sections). Following identification of specific gRNA(s), primers were designed by identifying regions containing gRNA target sites and finding primers that encompassed those regions (**Table S5**). The genomic DNA samples were then amplified and genotyped a second time using the primer set(s) specific to the target regions in the sample.

### Metabolic Analysis

All metabolic analyses were conducted using an Agilent SeahorseXF96 extracellular flux analyzer. Cell culture microplates (Agilent 102601-100) were seeded with 50,000 cells 24 hours prior to running the assay. Sensor cartridges (Agilent 102601-100) were hydrated with sterile water and incubated, along with XF calibrant (Agilent 100840-000), in a non-CO_2_ incubator 24 hours prior to use. Complete Seahorse assay medium (Agilent 103680-100) was made immediately prior to running the assay according to the manufacturer’s instructions. 160µL of XF calibrant was added to the entirety of the plate. The cell culture microplate and sensor cartridge were then incubated at 37°C in a non-CO_2_ incubator. All assays performed used the Seahorse XF Cell Mito Stress Test Kit (Agilent 103015-100) with Oligomycin 1.5µM, FCCP 1.0µM, Rotenone/Antimycin A 0.5µM, compounds were reconstituted and diluted using complete seahorse medium on the day of the assay. Cell number normalization was performed through image-based counting of cells prior to running the assay (using the InCell as described above).

## Supplementary Methods

### Cell Picking

Each raft has a four-character alphanumeric coordinate (Raft ID). Fiduciary markers located at fixed locations on the raft plate (shown below) in conjunction with custom software (FIVTools > Cal Check, Calibration) were used to locate individual rafts for map generation (see **Image Analysis and Quality Control**).

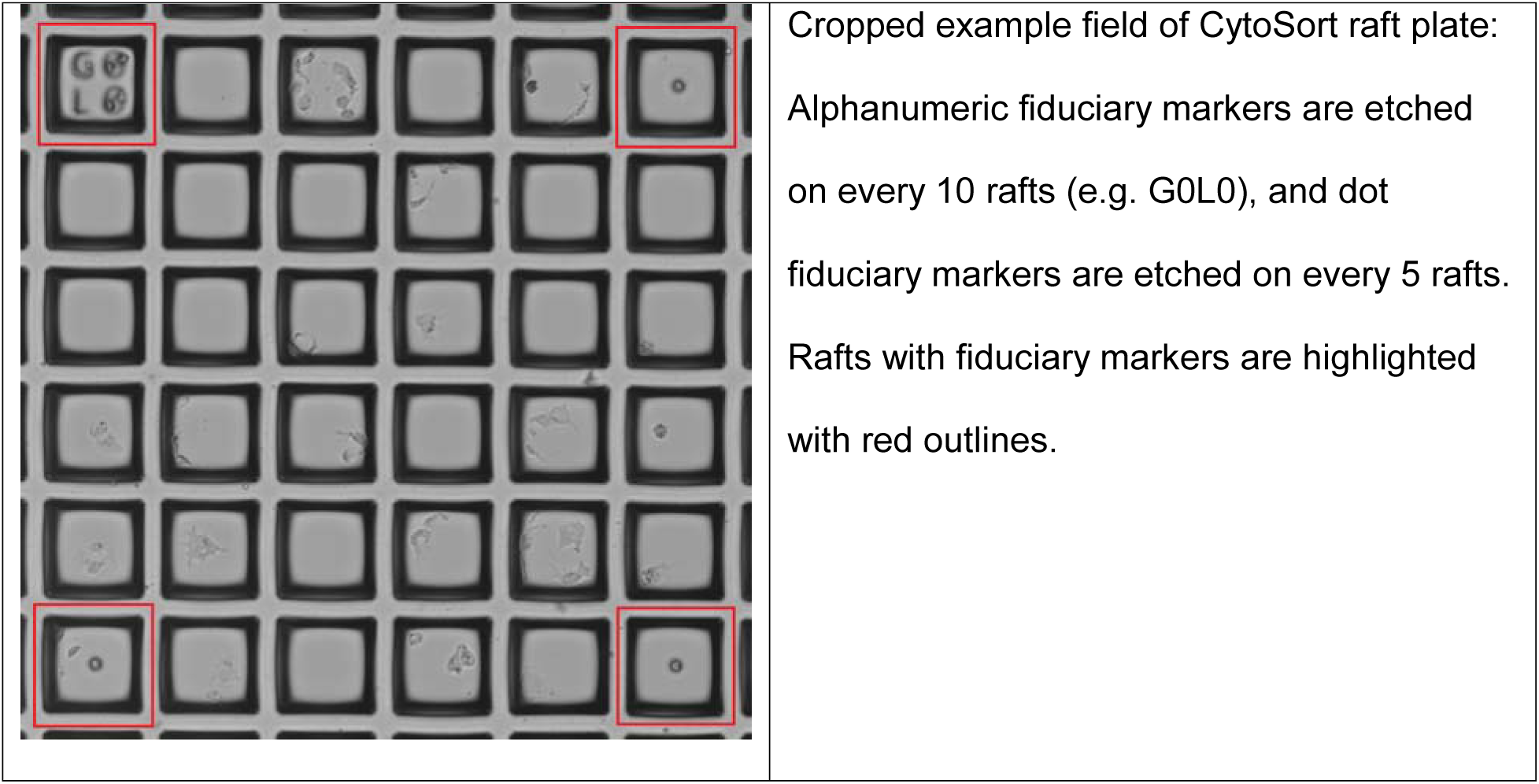

### HEK293 Cell Nucleofection

SF cell line solution stock was prepared from the Lonza SF Cell line 4D-Nucleofector LV Kit XL (V4LC-2520) kit by combining 82µL of SF Cell Line Nucleofector^TM^ Solution with 18µL Supplement 1. 20µL of stock solution was then combined with 2µL 1mg/mL Cas9 protein (QB3 MacroLab) and 2µL 1mg/mL gRNA. The complexes were incubated on the benchtop for 10 minutes. Cell suspensions of 2×10^5^ cells were placed in 1.5 mL microcentrifuge tubes and spun down. Supernatant was carefully removed avoiding the cell pellet. The cells were rinsed in 1x PBS and centrifuged a second time. Supernatant was carefully removed, and the cells were resuspended in the final combined SF cell line solution and transferred to a nucleofector cuvette, found in the nucleofector kit. The cuvette was placed in the Lonza 4D-Nucleofector Unit. (Lonza AAF-1002X, AAF-1002B) and nucleofected with Pulse code CM130. Nucleofected material was added to a prewarmed 6 well plate with 5mL of DMEM media in each well. Nucleofected cells were incubated for 48 hours for recovery. The cell line solution is not healthy for the cells, so speed is a priority upon resuspension in cell line solution.

### RFP-GFP Cell Perturbation and Isolation

A HEK293 cell line expressing both RFP and GFP (Gentarget #SC009) was used in this experiment. Cells were then nucleofected with a gRNA targeting GFP (TGCCCGAAGGCTACGTCCAG) or RFP (GGCCACGAGTTCGAGATCGA) and Cas9. All gRNAs were ordered from Synthego. Nucleofection was conducted using a Lonza 4D-Nucleofector Unit. (Lonza AAF-1002X, AAF-1002B). Following the Raft-Seq workflow, cells were imaged and the guide presence was predicted by a combination of RFP and GFP intensity features. Cells were selected and isolated into 96-well plates. Alternatively, cells were sorted using a Sony SH800S cell sorter individually by their RFP and GFP fluorescence. Cells were then genotyped and designated as being given the RFP or GFP guide if the locus that the respective guide targets had been altered.

### Modeling Considerations

We implement several specific design criteria during the experimental setup and model selection. First, the ‘pure’ samples are split into testing wells and training wells so that we can decrease overfitting during model selection. For most of the experiments presented herein, our training datasets comprised two of the three labeled population wells of each type—the two wells must come from different plates—and use the remaining two labeled wells as “testing sets”. We split the data by well rather than randomly assigning cells from all pure population wells to mitigate the influence of batch effects during modeling. It is important to note that the actual data used for ultimate validation of the models will be the identities of the perturbagen in the unlabeled cell population. These are independent of the model scores of the pure populations, so we do not have to be concerned about hyperparameter overfitting from examining the testing data. Second, the testing wells are spread across different plates to hopefully account for batch effects. Since training data and testing data technically come from separate samples, we observed several odd phenomena such as prediction quality improving for testing data. These phenomena would most likely not be observed if the model-generating data were truly randomized, but we felt it was important to retain any variance caused by batch effects, since batch effects present a large and consistent obstacle to cell imaging analysis (Caicedo et al.).

Relatedly, we make sure not to create any artificial batch effects generated by inconsistencies in the screening process. All stains used are prepared as a single batch to be used across plates. All imaging and feature extraction settings are kept constant across each plate in a screening experiment.

Another aspect of model design that we consider is the exclusion of ‘leaky’ variables, or variables that happen to correlate with the data labels. Since our labeled data exists on specific positions of the plate, including a variable that is effectively a proxy for cell position would render the model useless on unlabeled data. We have found that several variables that are in the standard output of IN Carta feature extraction are leaky, such as features measured on a global basis, and we exclude them before starting the modeling process.

We also face a time constraint in our modeling pipeline. This time constraint is caused by the necessity of the locations of cells on the plate staying constant between imaging and isolation. Though we can fix the cells, a modeling pipeline based on that assumption would limit our ability to perform Raft-Seq for the isolation of live cells. Optimally, we complete feature selection, modeling, and raft selection within six hours of imaging, and continue on to isolation. This of course, requires a lot of computational power.

## Supplementary Data

**Figure S1:**
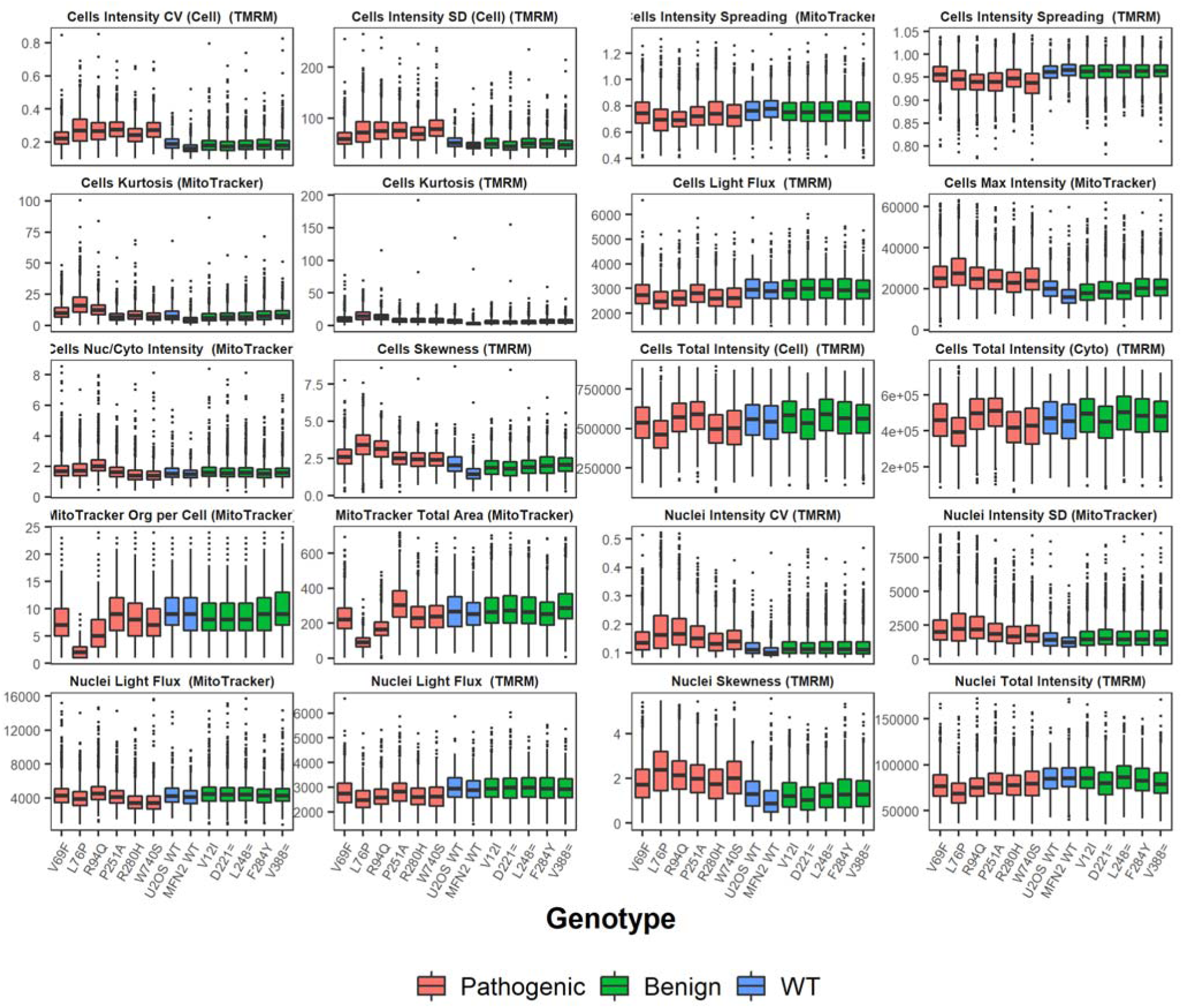
Box plots of important features extracted from TMRM and MitoTracker staining. Histograms are colored by whether the cell was wild type (unmodified wild type or modified with a wild type MFN2 cDNA), contained MFN2 cDNA containing a benign mutant, or contained MFN2 cDNA containing a pathogenic mutant. Features that showed no apparent difference between mutants were omitted.

**Figure S2:**
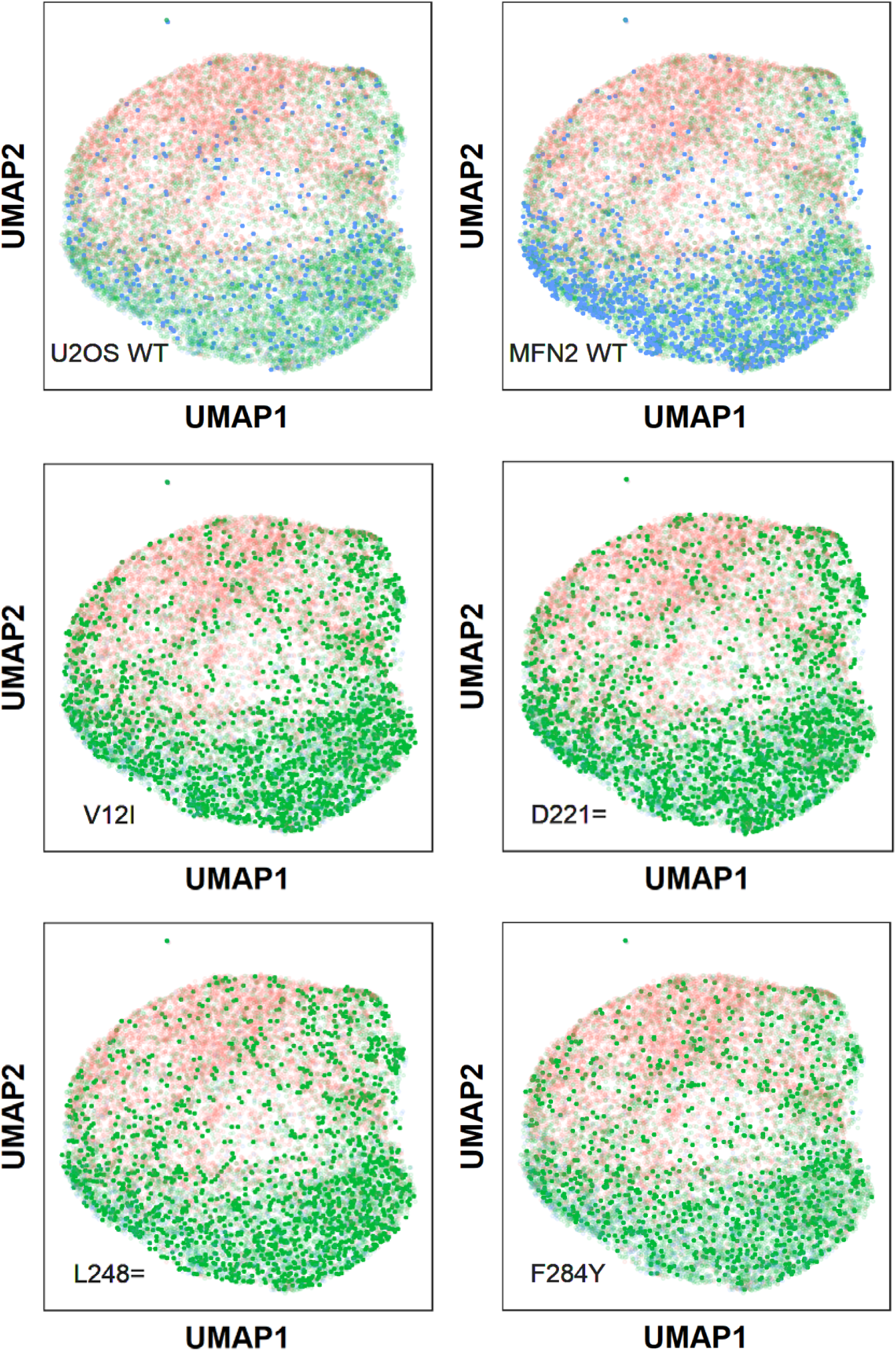
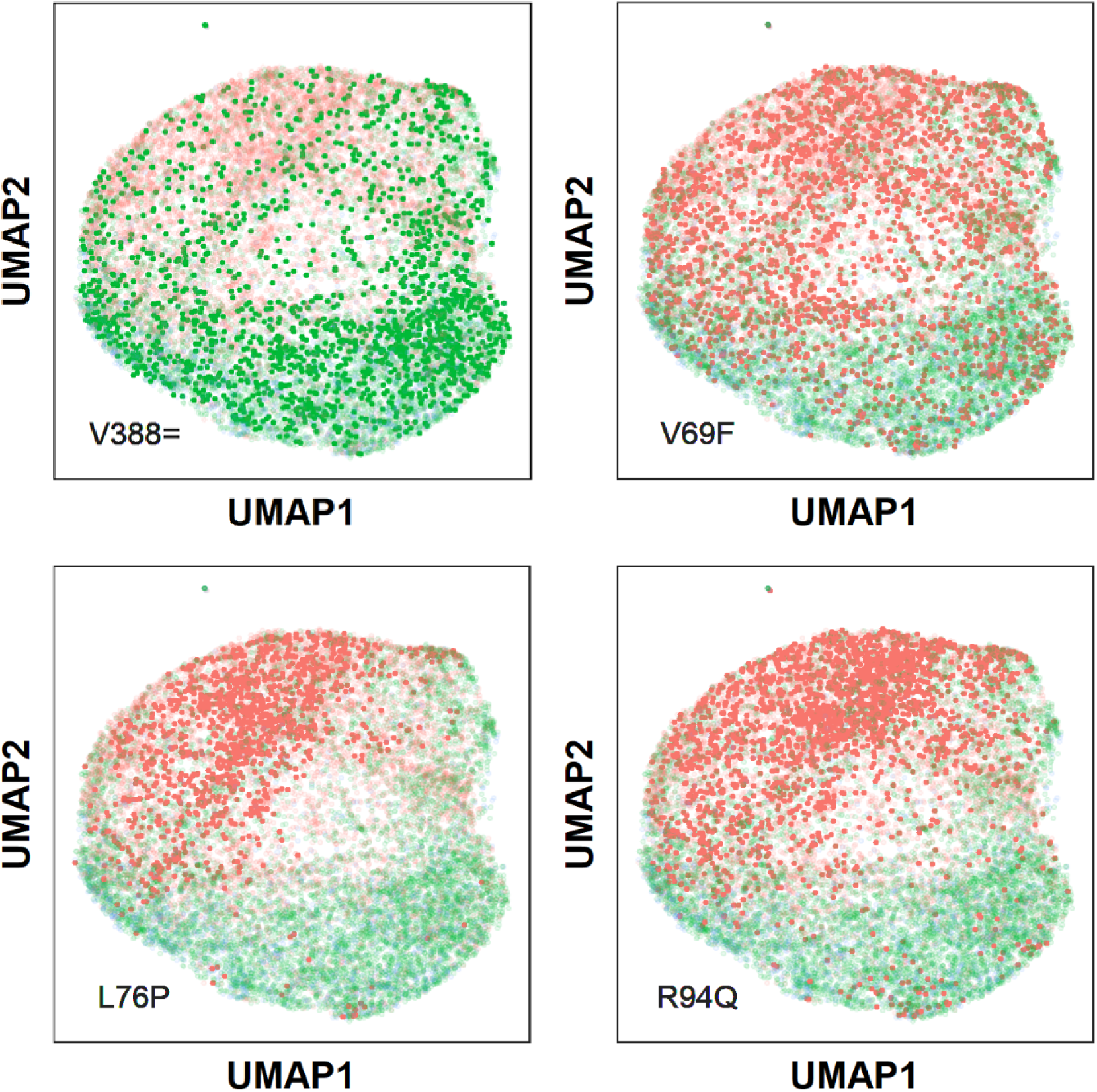
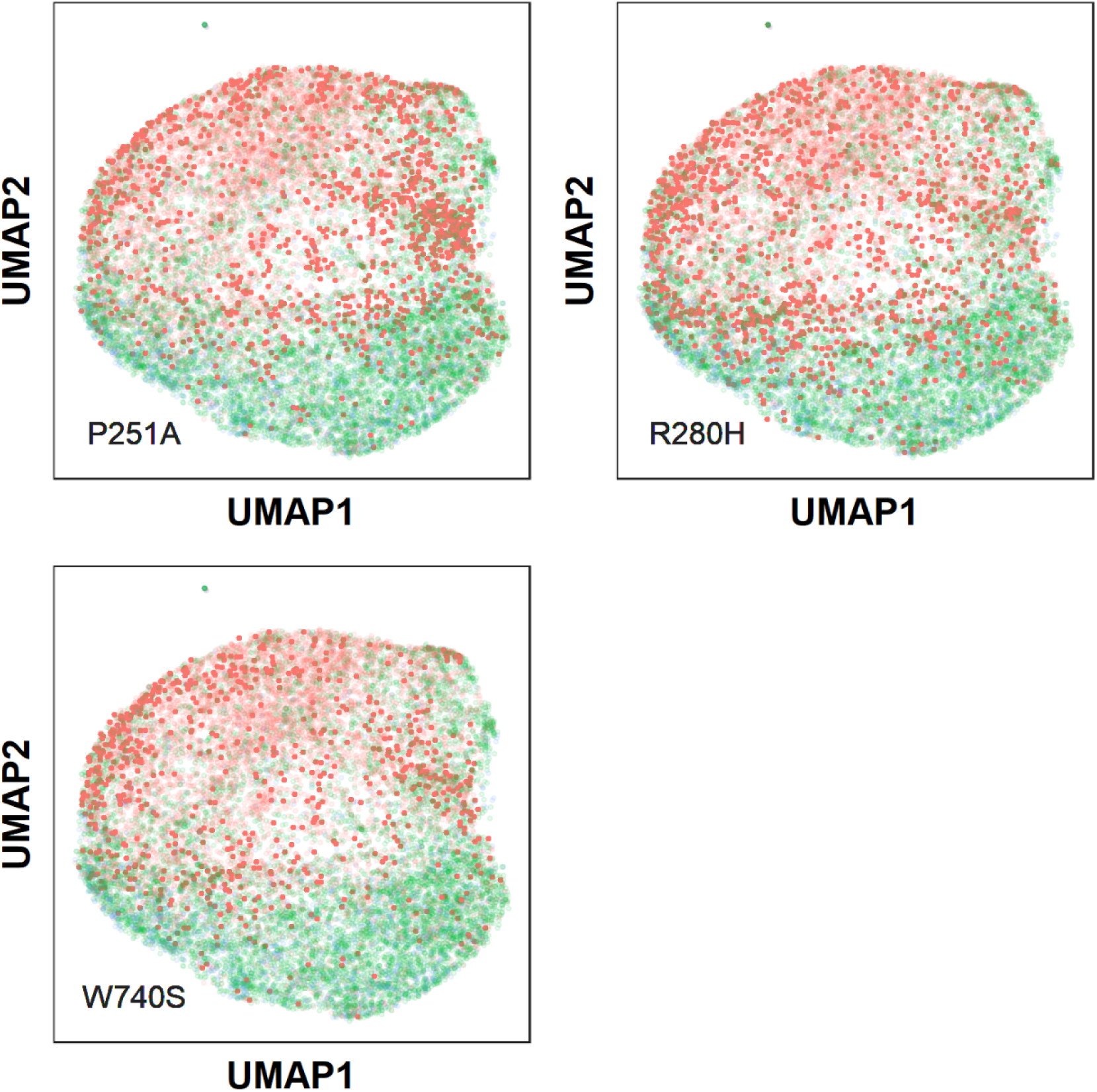
Scatter plot for cell feature data UMAPs, highlighted by cell genotype. In *blue*, U2OS WT refers to cells uninfected with lentivirus, while *MFN2* WT refers to cells infected with lentivirus containing the wild type *MFN2* gene. Otherwise, the highlighted genotype refers to the mutation present in the introduced *MFN2* gene (*red* pathogenic, *green* benign).

**Figure S3.**
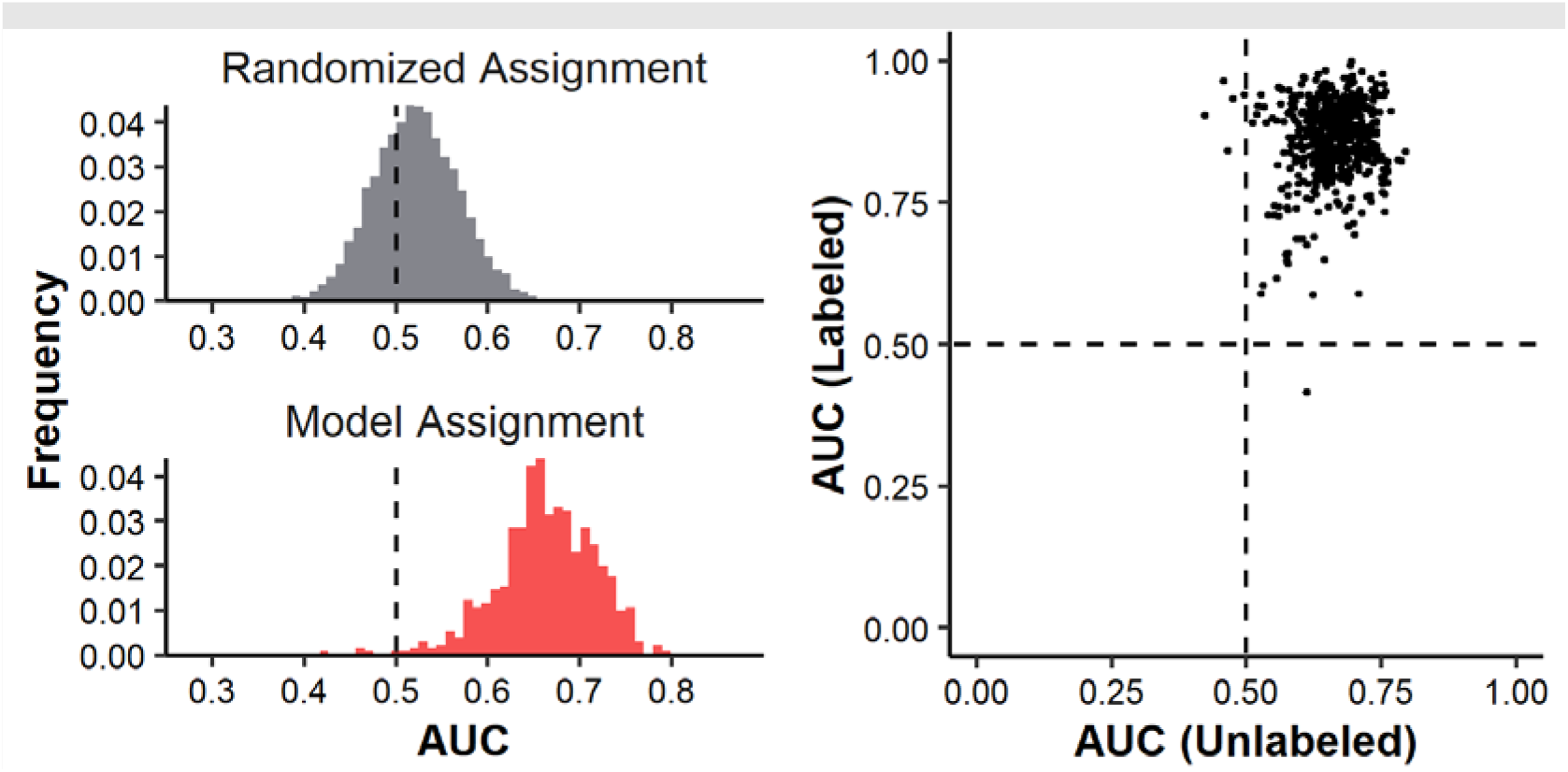
Model AUCs for identifying a mixture of wild type and six pathogenic mutant cell lines: **Left**: A histogram of AUCs for models detecting mutants in the unlabeled population of wild-type cells and mutants. On top, a histogram of AUCs generated from randomly assigning models is shown as a comparison. **Right**: A scatterplot showing the performance of 610 models in detecting mutants in a mixture of wild type cells and four pathogenic mutants. Each point represents a model and its position is determined by its ability to distinguish cells in the pure control wells and cells in the mixed wells.

**Figure S4.**
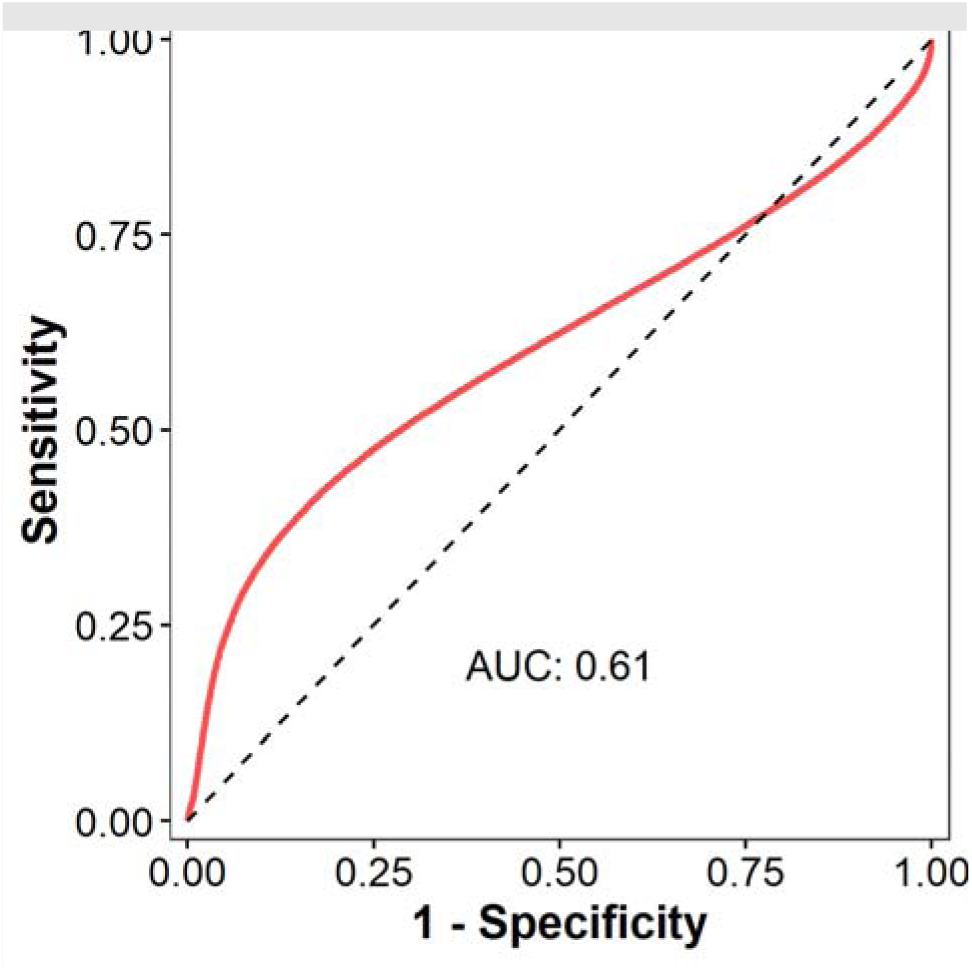
Modeled ROC from flow cytometry: ROC Curve based on flow data for *MFN2* WT vs. *MFN2* R280H mutants on the PerCP-Cy5.5-A-Compensated channel.

**Figure S5.**
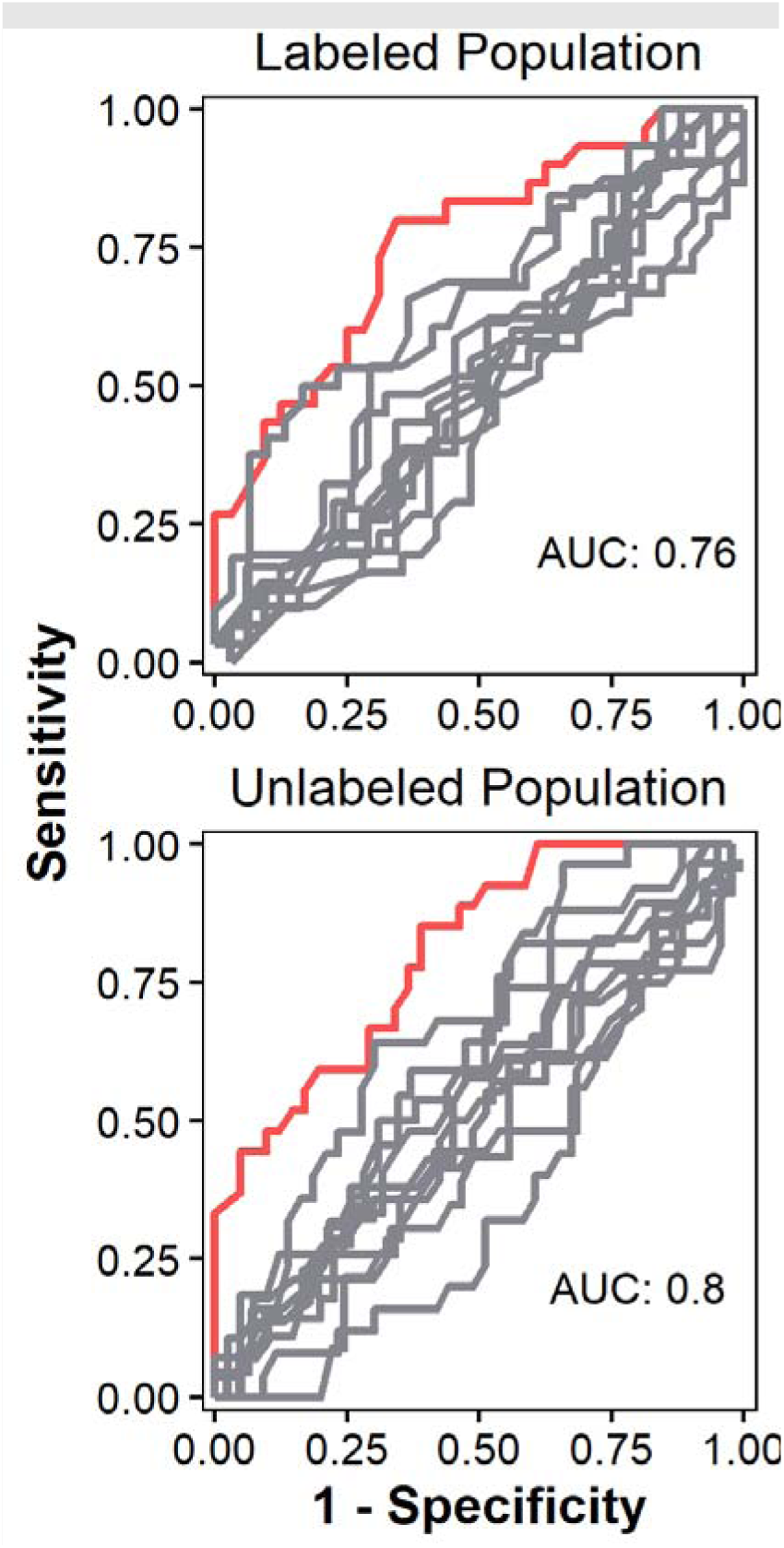
Results from Raft-Seq between P251A Pathogenic and D221= Benign: ROC Curves which were generated using the best model, separated by data that was generated from cells picked from wells in the labeled cell population (upper) and from the unlabeled cell population (lower). The red curves are for the models in question and the grey curves are those for control models generated by random shuffling of labels.

**Figure S6.**
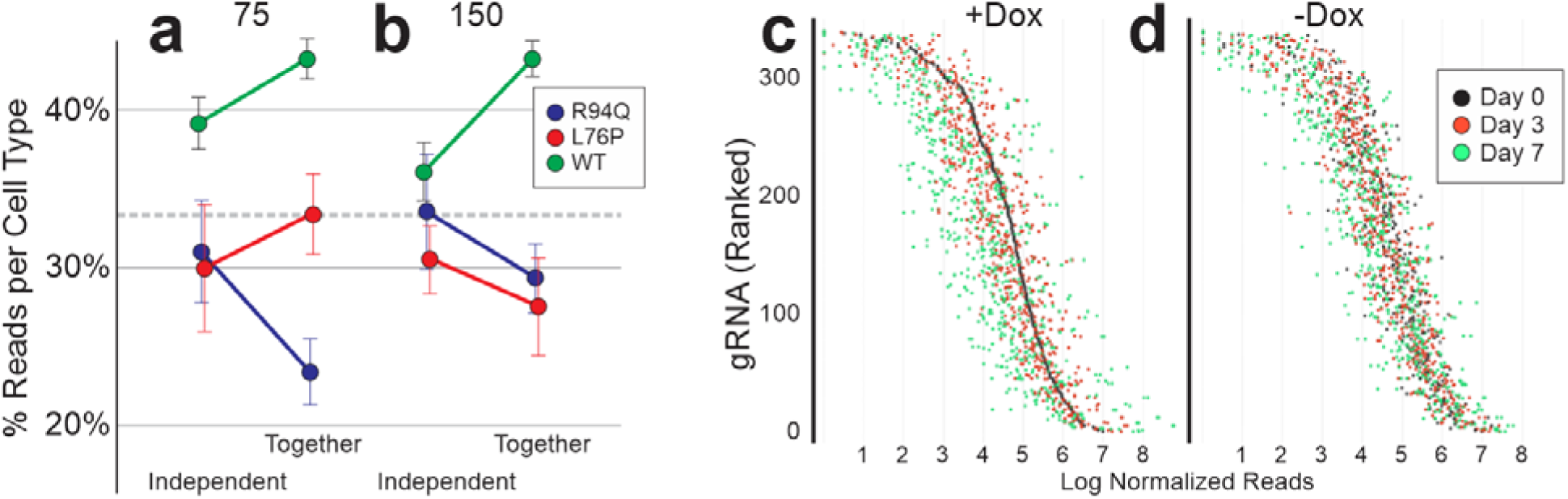
Fitness consequences of MFN2 mutations. **a,b**) U2OS stable lines containing R94Q, L76P, mutations, or the WT control line were grown in separate wells or evenly mixed together. This was done at 2 densities 75,000 (**a**) and 150,000 (**b**) cells / well of a 6-well plate and cells were allowed to grow for 4 days. Cells grown independently were harvested into the same tube (so all subsequent steps would be together). All cells were taken through the genotyping pipeline using the same primers as above, then the reads were counted, and the proportion of each cell type measured. **c,d**) A similar experiment was performed with the scanning MFN2-targeted gRNAs, under Dox (**c**) to induce Cas9 cutting, or under control conditions (**d**) with the gRNAs, but no Dox (and therefore no Cas9 cutting). Samples were taken initially, then at 3 and 7 days after Dox induction. Bulk timepoint samples were processed through the genotyping pipeline and reads were counted for each gRNA.

**Figure S7.**
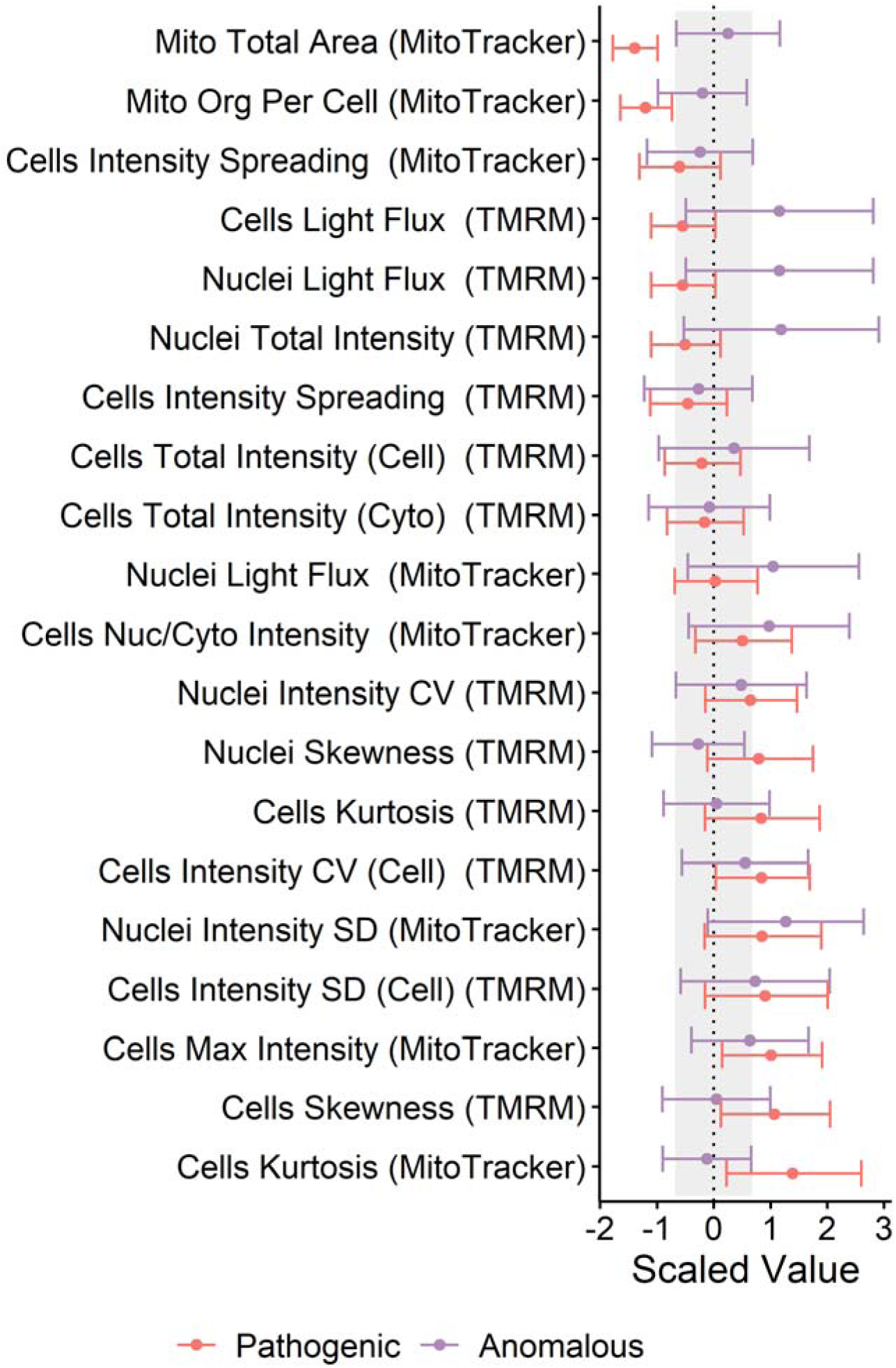
Isogenic Feature Comparison. We compared the distributions of the important features identified earlier for anomalous cells and strong pathogenic mutants (L76P & R94Q) by normalizing each to the distribution of the non-anomalous and benign mutants, respectively. Some of the features covaried between pathogenic and anomalous, and this pattern was stronger than in the non-anomalous (pathogenic vs. non-anomalous r^2^ -0.02, vs anomalous r^2^ 0.001, neither significant). Red is for the distribution in the pathogenic cells; purple is for the anomalous cells. Background grey shading indicates the base distribution for the non-anomalous/benign cells. Bar width and ranges correspond to 0.687σ in both directions, covering 50% of the data. Anomalous cells are the top 0.5% highest anomaly-detection scores from 152 models.

**Figure S8:**
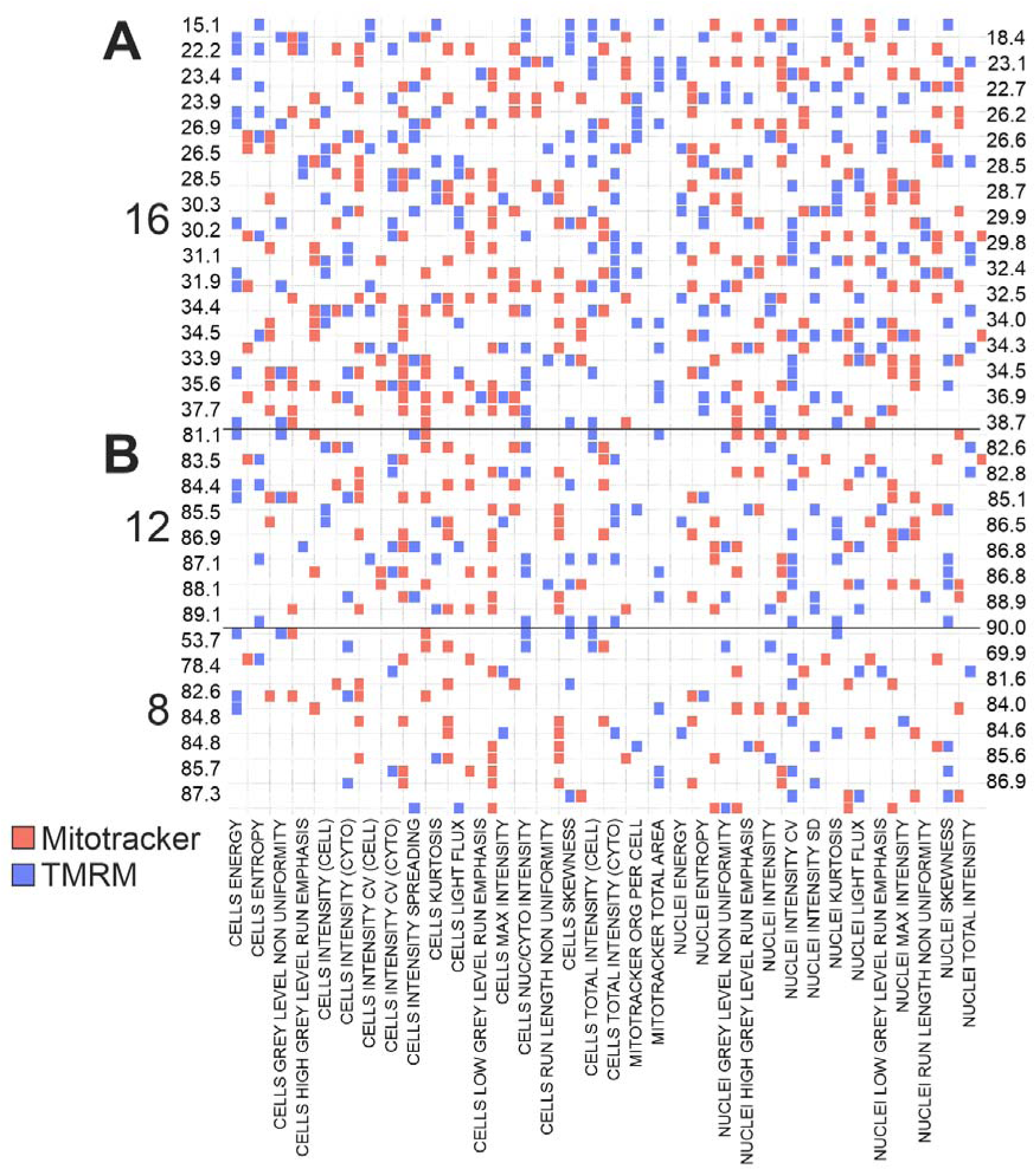
MFN2 Isogenic Lines carrying mutations are distinct from WT. Decision Jungles were trained on either WT vs. Known KO clones (**A**) or known mutants and known un-cut clones (**B**) with either 16, 12, or 8 features per model. Each model had a random set of features it could use in training. Then, the model was deployed and the Kruskall-Wallis p-Value comparing the model scores of the UTR/Coding mutants with the WT clones was assessed (these p-values are displayed as -log10 on either side of the graph).

**Figure S9.**
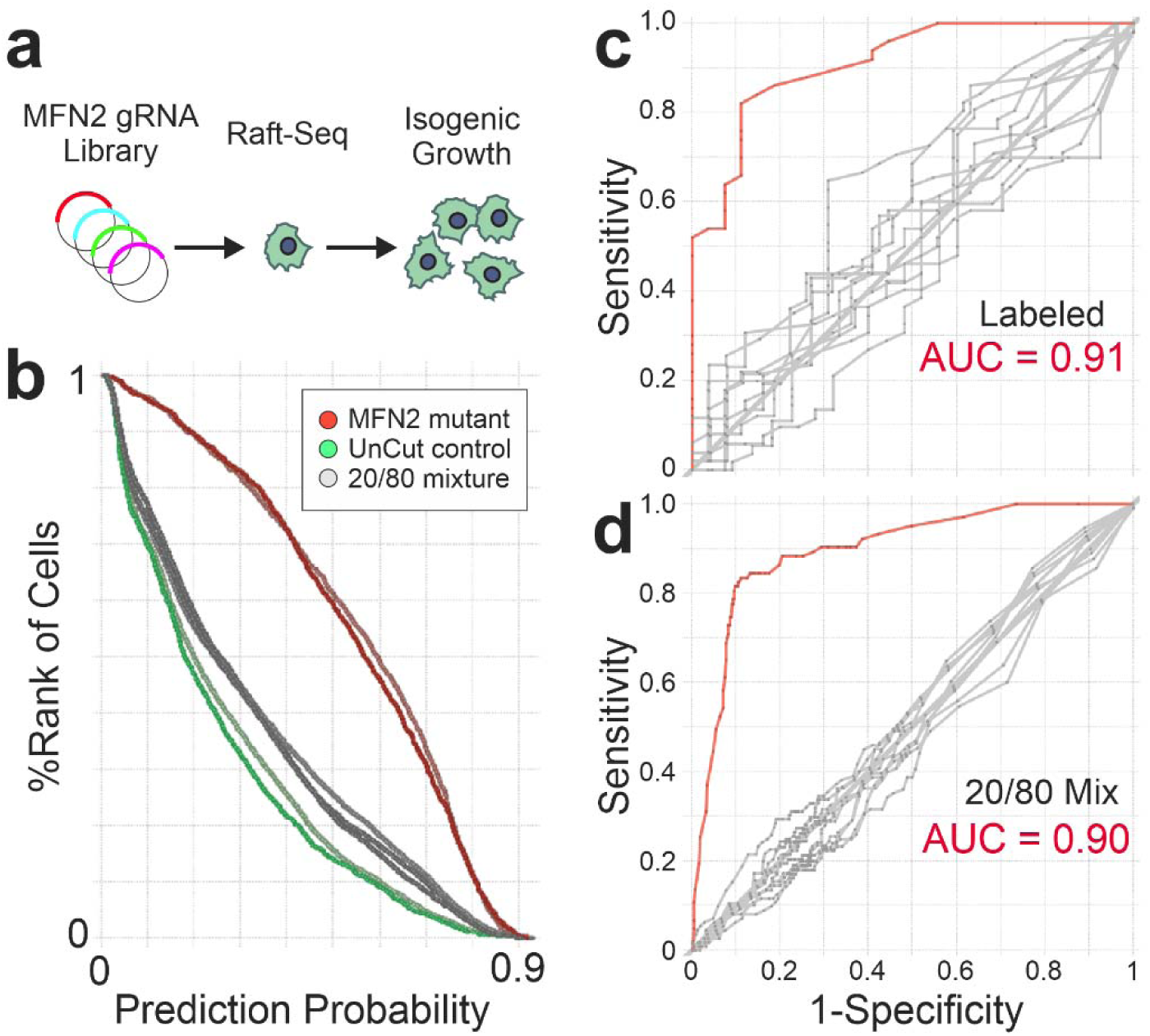
Endogenous MFN2 Mutant Line Can Be Effectively Separated from a Non-Cutting Control. **a**) U2OS cells with were screened against the MFN2 gRNA library using Raft-Seq, and single cells were picked for isogenic growth. One line, 5’ UTR Ins A04, was used here in a Raft-Seq experiment to compare its ability to separate against an un-cut control. The model was trained in AzureMLStudio (two class decision jungle) with 5 features (Nuclei Area, Mitotracker Intensity Kurtosis, Skewness, Energy and TMRM Intensity Entropy). **b**) Ranked histogram comparing single cells from labeled wells (where only the mutant or control were plated) against their prediction probabilities from the 5-feature model. Cells from the control (labeled) wells, and mixture well (20% mutant, 80% control) were picked and genotyped. **c**) ROC curves for the labeled (control) well (n=77). **d**) ROC curve from the wells with a mixture of the two cell types. The model performed with an AUC = 0.9, accuracy of 88%, precision 69% (n = 490 cells). Red curves are the real data while the gray curves were generated from sets of data where the link between the true genotype and the prediction probability were scrambled (noise models).

**Table S1:**
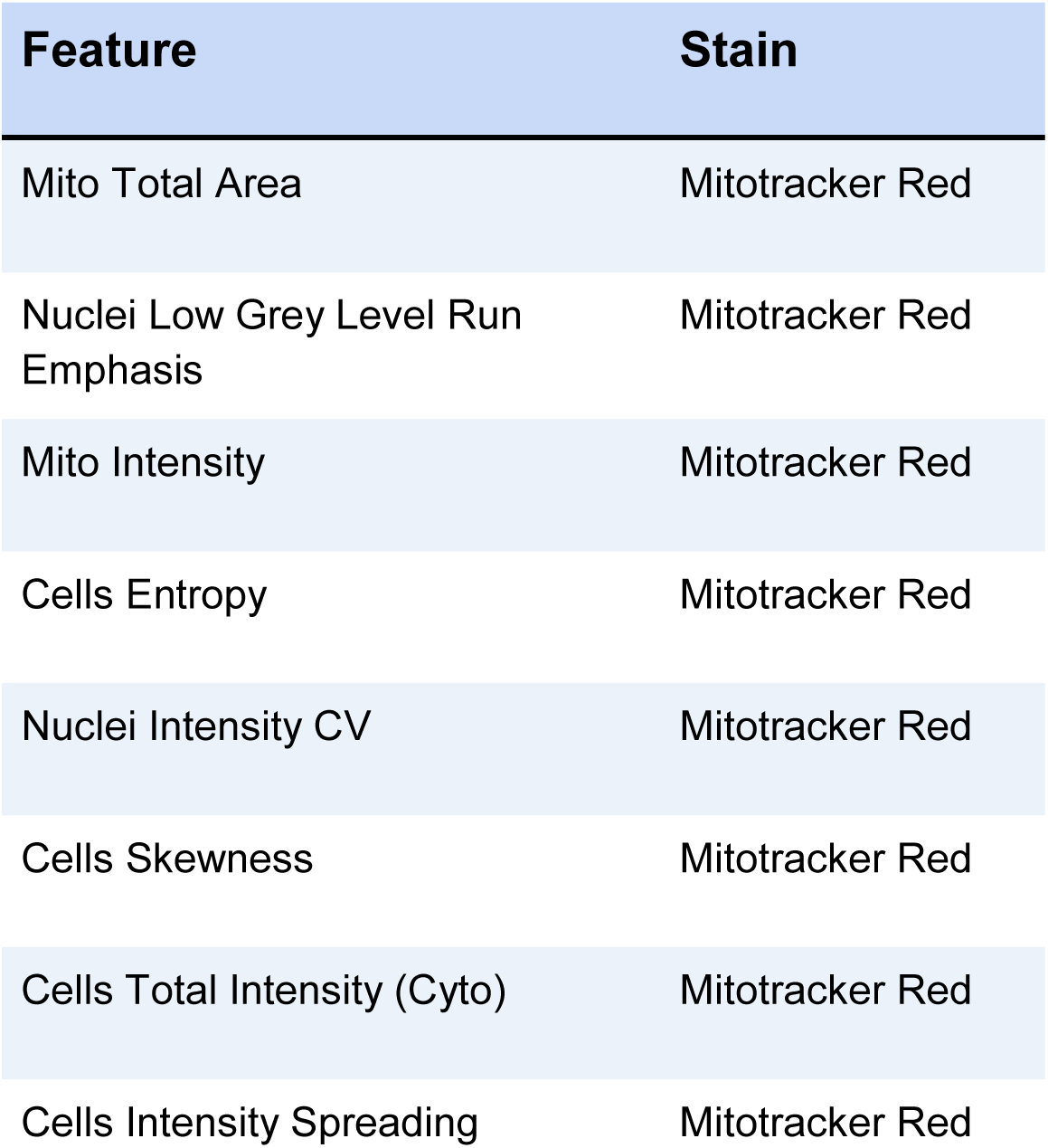
Table of the 8 features used for cell isolation and random feature assembly with the corresponding stain.

**Table S2:**
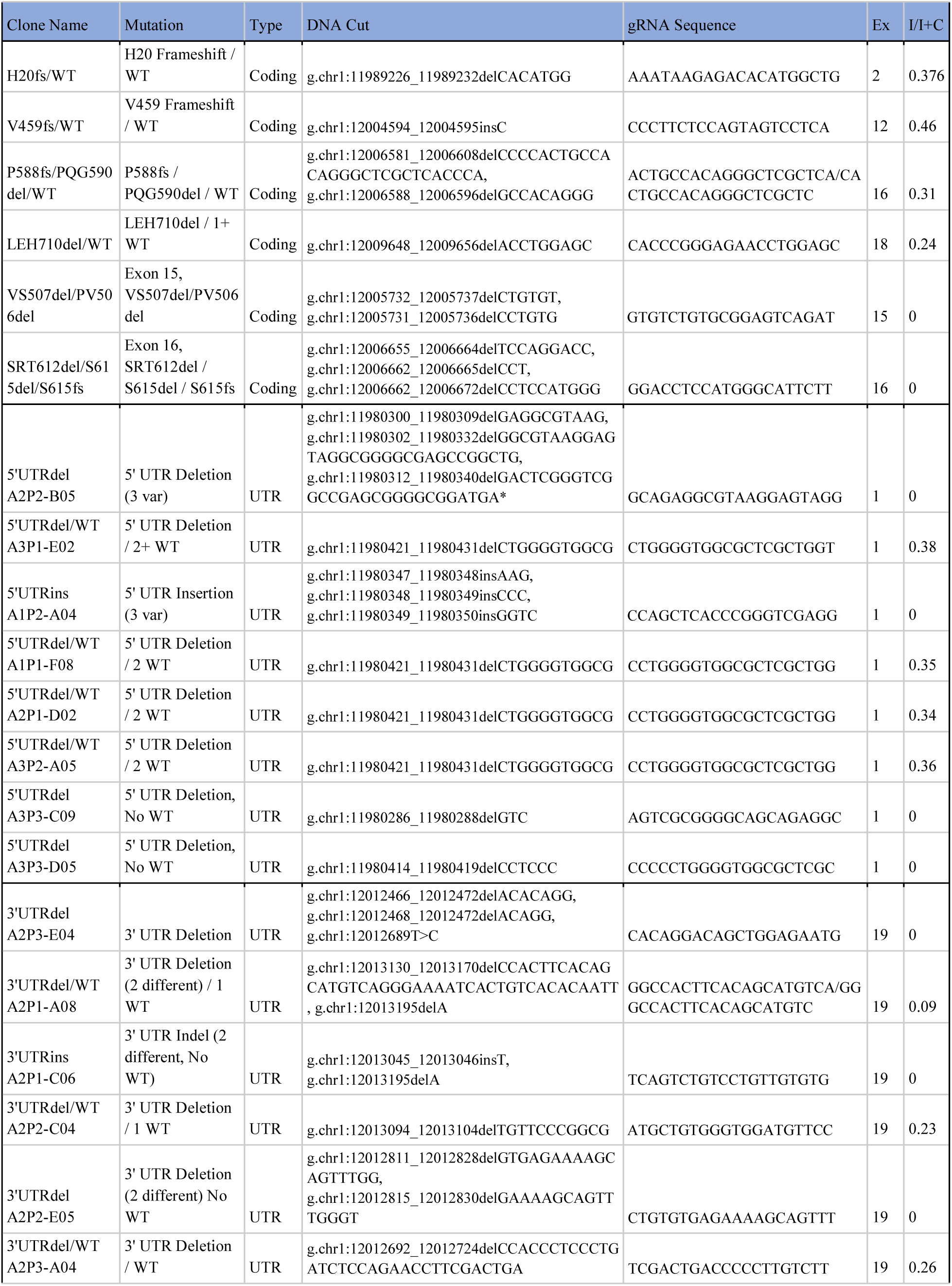
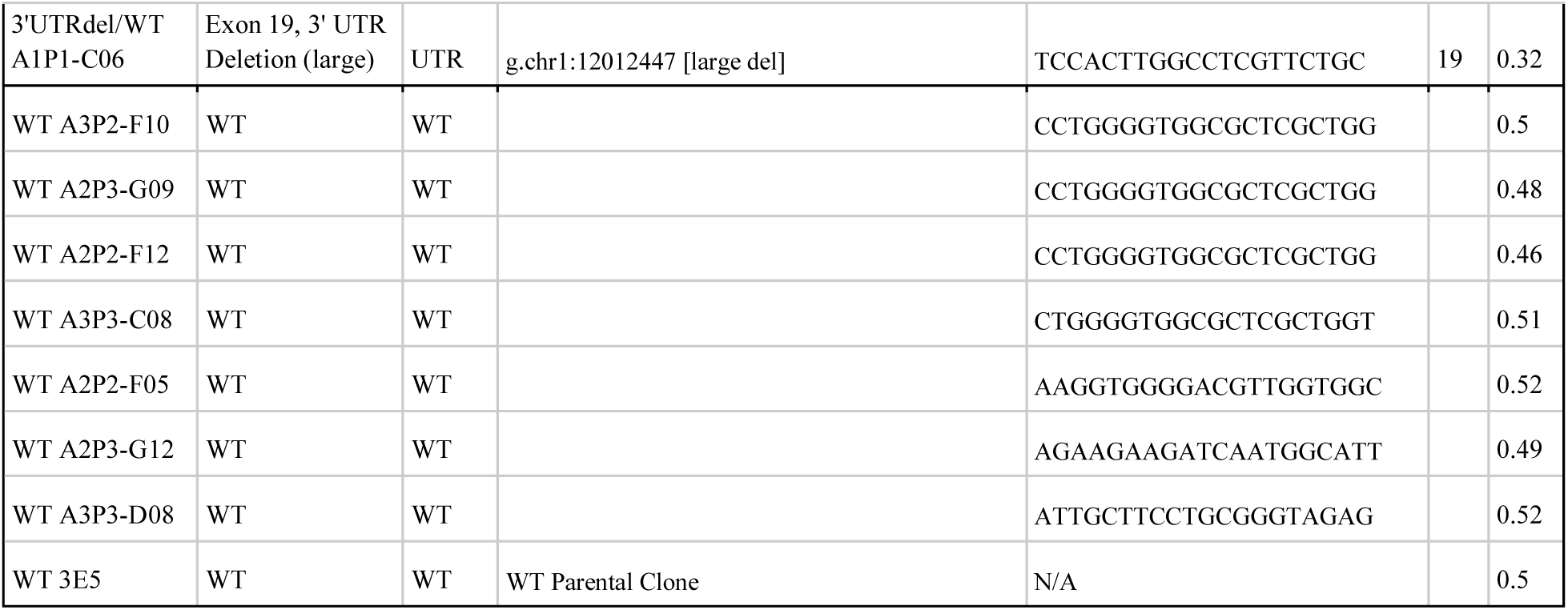
*MFN2* Isogenic mutant line information table. The U2OS isogenic lines were genotyped to determine the gRNA and the editing consequence of Cas9 cutting. The table shows the ‘Clone Name’ used in other parts of the paper as well as the longer mutation definition for the clone and the specific genomic mutation. Also listed is the sequence of the gRNA used, the exon where the cut was located, and the % of intact target reads as a ratio with a control (constant) sequence, where 0 = all alleles are mutant, and 0.5 = all alleles are WT. Variant calling was performed after paired end reads were merged using the fastq-join method from ea-tools (https://github.com/ExpressionAnalysis/ea-utils). Merged fastq files were then aligned using BWA-mem (DOI:10.1093/bioinformatics/btp324). Annotated variant call files were produced using Picard Tools (http://broadinstitute.github.io/picard/) and the Genome Analysis Tool Kit (DOI: 10.1101/201178).

**Table S3:** Individual mitochondrial features that distinguish the isogenic mutants from un-cut clones. The name of the engineered feature and the Kruskal-Wallis p-Value distinguishing the single feature’s mean difference between the cut isogenic clones (UTR or coding), and the WT isogenic clones. WVM refers to Mitotracker and WVT refers to TMRM. Features with Level in the name are texture-based features.

| Mitochondrial Feature | KW p-value |
| --- | --- |
| CELLS ENTROPY WVT | 2.72E-17 |
| CELLS ENTROPY WVM | 4.40E-16 |
| CELLS KURTOSIS WVT | 2.62E-15 |
| CELLS ENERGY WVT | 8.06E-15 |
| CELLS KURTOSIS WVM | 2.06E-13 |
| CELLS SKEWNESS WVM | 3.11E-13 |
| CELLS INTENSITY (CYTO) WVM | 3.10E-12 |
| CELLS ENERGY WVM | 1.59E-10 |
| CELLS INTENSITY (CELL) WVM | 1.64E-10 |
| CELLS INTENSITY (CYTO) WVT | 2.42E-10 |
| CELLS GREY LEVEL NON UNIFORMITY WVM | 2.60E-10 |
| CELLS RUN LENGTH NON UNIFORMITY WVT | 3.41E-10 |
| NUCLEI ENTROPY WVT | 4.33E-10 |
| CELLS RUN LENGTH NON UNIFORMITY WVM | 4.78E-10 |
| CELLS LOW GREY LEVEL RUN EMPHASIS WVT | 7.24E-10 |
| CELLS INTENSITY (CELL) WVT | 2.49E-09 |
| CELLS SKEWNESS WVT | 6.26E-09 |
| NUCLEI ENERGY WVT | 9.21E-09 |
| CELLS TOTAL INTENSITY (CYTO) WVT | 2.23E-08 |
| CELLS TOTAL INTENSITY (CELL) WVT | 1.21E-07 |
| NUCLEI ENTROPY WVM | 2.66E-07 |
| CELLS TOTAL INTENSITY (CYTO) WVM | 9.80E-07 |
| CELLS TOTAL INTENSITY (CELL) WVM | 3.76E-06 |
| NUCLEI LOW GREY LEVEL RUN EMPHASIS WVT | 7.18E-06 |
| CELLS GREY LEVEL NON UNIFORMITY WVT | 8.21E-06 |
| NUCLEI RUN LENGTH NON UNIFORMITY WVT | 2.34E-05 |
| CELLS MAX INTENSITY WVT | 3.08E-05 |
| NUCLEI GREY LEVEL NON UNIFORMITY WVM | 5.76E-05 |
| NUCLEI TOTAL INTENSITY WVT | 1.00E-04 |
| NUCLEI SKEWNESS WVM | 2.04E-04 |
| CELLS LOW GREY LEVEL RUN EMPHASIS WVM | 4.76E-04 |
| NUCLEI INTENSITY WVT | 5.42E-04 |
| NUCLEI TOTAL INTENSITY WVM | 5.72E-04 |
| NUCLEI MAX INTENSITY WVT | 5.93E-04 |
| NUCLEI INTENSITY WVM | 8.27E-04 |

**Table S4:** Metabolic testing results on the U2OS *MFN2* mutant isogenic cell lines. The ‘Clone Name’ for each isogenic line is given, the average and upper/lower 95% confidence intervals (in parentheses) of the OCR (pMoles/min) normalized to parental WT control (iCas9-3E5). The 4 columns indicate baseline OCR vs the OCR after adding the named toxins. Mutant clones are sorted so that normal basal activity is listed first with more disrupted activity listed further down, row borders are used as a visual aid only.

| Clone | Baseline | FCCP | Oligomycin | ROT/AA |
| --- | --- | --- | --- | --- |
| iCas9-3E5 | 1 (0.99/1) | 2.02 (2/2.04) | 0.39 (0.38/0.39) | 0.27 (0.27/0.27) |
| WT A2P2-F12 | 1.02 (1.01/1.03) | 1.5 (1.43/1.58) | 0.44 (0.42/0.46) | 0.25 (0.24/0.25) |
| WT A2P3-G12 | 0.8 (0.79/0.81) | 1.69 (1.66/1.72) | 0.34 (0.34/0.34) | 0.26 (0.26/0.27) |
| WT A3P3-C08 | 0.77 (0.77/0.78) | 1.52 (1.46/1.58) | 0.27 (0.27/0.27) | 0.21 (0.21/0.21) |
| WT A3P3-D08 | 1.02 (1.01/1.02) | 1.86 (1.78/1.95) | 0.42 (0.41/0.43) | 0.29 (0.29/0.3) |
| 3'UTRdel/WT A2P3-A04 | 0.97 (0.95/0.98) | 2.02 (2/2.04) | 0.32 (0.32/0.32) | 0.21 (0.21/0.22) |
| 5'UTRdel A2P2-B05 | 0.95 (0.93/0.97) | 2.11 (1.97/2.25) | 0.33 (0.33/0.33) | 0.22 (0.21/0.22) |
| 5'UTRins A1P2 | 0.88 (0.88/0.89) | 1.77 (1.74/1.8) | 0.34 (0.34/0.34) | 0.25 (0.25/0.25) |
| 3'UTRdel A2P3-E04 | 0.87 (0.86/0.88) | 1.99 (1.98/2.01) | 0.34 (0.34/0.35) | 0.2 (0.2/0.21) |
| 3'UTRdel/WT A2P1-A08 | 0.74 (0.73/0.76) | 1.48 (1.45/1.5) | 0.29 (0.28/0.29) | 0.19 (0.19/0.2) |
| 3'UTRdel A2P2-E05 | 0.72 (0.71/0.72) | 1.6 (1.58/1.62) | 0.26 (0.25/0.26) | 0.19 (0.18/0.19) |
| MFN2 L76P | 0.7 (0.7/0.71) | 1.2 (1.18/1.21) | 0.28 (0.28/0.28) | 0.2 (0.2/0.2) |
| H20fs/WT | 0.68 (0.67/0.68) | 1.29 (1.28/1.31) | 0.28 (0.28/0.28) | 0.21 (0.21/0.21) |
| V459fs/WT | 0.6 (0.59/0.61) | 1.05 (1.01/1.09) | 0.22 (0.21/0.22) | 0.15 (0.15/0.15) |
| VS507del/PV506del | 0.56 (0.55/0.58) | 1 (0.97/1.03) | 0.23 (0.23/0.24) | 0.17 (0.17/0.17) |
| 3'UTRdel/WT A2P2-C04 | 0.55 (0.54/0.56) | 1.07 (1.05/1.08) | 0.18 (0.18/0.18) | 0.11 (0.11/0.11) |
| 5'UTRdel A3P3-C09 | 0.44 (0.43/0.45) | 0.77 (0.73/0.8) | 0.15 (0.15/0.16) | 0.09 (0.09/0.1) |
| SRT612del/S615del/S615fs | 0.31 (0.31/0.32) | 0.53 (0.51/0.54) | 0.14 (0.14/0.14) | 0.1 (0.1/0.1) |
| 3'UTRdel/WT A1P1-C06 | 0.2 (0.2/0.2) | 0.27 (0.26/0.28) | 0.09 (0.09/0.09) | 0.06 (0.05/0.06) |

**Table S5: Separate Excel File.**

## Definitions

MAVE: multiplexed assays of variant effects
DMS: deep mutational scanning
VUS: Variant of Uncertain Significance
Genotype: The specific point mutation present (R280H, WT, H20fs, etc)
Genotype class: WT, Benign, or Pathogenic (aka what is designated by the model)
Labeled Cell Population: Used for training the models
Unlabeled Cell Population: admixtures of labeled cells (like benign + pathogenic) or in screening, where the researchers are completely blind to the identity of individual cells
ROC: Receiver Operator Curve
AUC: Area under the curve.

## Acknowledgements

We would like to thank Kate Matsunaga, Will Lee, and Silas Kuang for their help in the laboratory. We would also like to thank the Milbrandt lab for their continued support and equipment use. Genome Engineering and iPSC Center at Washington University School of Medicine also provides invaluable support to all members of our laboratory. We also thank the Vindigni lab for allowing us to utilize the engineered lines. Finally, we would like to thank the genetics administration and our cleaning staff.

## Author Contributions

ALY, JCB, and WJB wrote the manuscript. WJB and JCB designed the experiments. Initial planning and ideas brought by WJB, JDM, RDM. JEW, CLK, MJL, XEX, MAV, and ALY performed the experiments. JEW, CLK, MJL, XEX, JCB, ALY, VC, and WJB analyzed results. VC wrote some of the key software. All reviewed the paper and gave suggestions.

